# A Computational Model of Thalamic Tonic-burst Motor Projection Neurons and Their Regulation by Pallidal GABAergic Neurons

**DOI:** 10.1101/806653

**Authors:** Md Ali Azam, Deepak Kumbhare, Ravi Hadimani, Jamie Toms, Mark S. Baron, Jayasimha Atulasimha

## Abstract

A modified computational model of pallidal receiving ventral oral posterior (Vop) thalamocortical motor relay neurons was adapted based on in vivo observations in our rodent model. The model accounts for different input neuronal firing patterns in the primary motor output nucleus of basal ganglia, the globus pallidus interna (GPi) and subsequently generate Vop outputs as observed in vivo under different conditions. Hyperpolarizing input de-inactivates its T-type calcium channel and sets the thalamic neurons in the preferable burst firing mode over a tonic mode and induces low threshold spikes (LTS). In the hyperpolarized state, both spontaneously and in response to excitatory (e.g. corticothalamic) inputs, burst spiking occurs on the crest of the LTS. By selecting and determining the timing and extent of opening of thalamic T-type calcium channels via GABAergic hyperpolarizing input, the GPi precisely regulates Vop-cortical burst motor signaling. Different combinations of tonic, burst, irregular tonic and irregular burst inputs from GPi were used to verify our model. In vivo data obtained from recordings in the entopedunucular nucleus (EP; rodent equivalent of GPi) from resting head restrained healthy and dystonic rats were used to simulate the influences of different inputs from GPi. In all cases, GPi neuronal firing patterns are demonstrated to act as a firing mode selector for thalamic Vop neurons.

## 1. Introduction

The basal ganglia (BG) processes a wide range of distinct cortico-motor signals in parallel and relays these modulated signals to the motor cortex via the thalamus (Chakravarthy et al. 2010; Parent and Hazrati 1995; Lanciego et al. 2012; Middleton and Strick 2000). The pallidal receiving ventral oral posterior (Vop) thalamic nucleus is the major relay nucleus for motor output from the BG to the motor cortices (Aldes 1988). The Vop relay neurons receives GABAergic input (Albin et al. 1989; Bolam et al. 2000) from the globus pallidus interna (GPi), the major motor output nucleus of BG, and transmits excitatory signals to the cortex (Parent and Hazrati 1995; Lanciego et al. 2012). Despite its importance, the mechanism by which the GPi regulates the thalamocortical drive has not been satisfactorily scrutinized. Moreover, little is known about how the basal ganglia ultimately influences motor behavior and such aspects as learning. Although the classical BG rate model (Albin et al. 1989; DeLong 1990) suggests that the inhibitory output from GPi acts to proportionally inhibit and thereby regulate thalamocortical drive, the model cannot, for example, account for the benefits of a GPi pallidotomy or deep brain stimulation (DBS) on *both* hypo- and hyperkinetic motor features in movement disorders. According to the rate model, reduced GABAergic resting activity in GPi should disinhibit (activate) thalamocortical motor drive. Moreover, also in conflict with the classical model, select phasic *increases* in neuronal discharge activity in the entopeduncular nucleus (EP; rodent equivalent of GPi) heralds focused motor activation in normal rats, while excess, synchronized, wide-spread phasic increases in GPi neuronal activity heralds excess, widespread muscle contractions in dystonic rats (Kumbhare et al. 2015; Baron et al. 2011).

During in vivo recordings from healthy rats under head restrained resting conditions, we consistently observed that, when the majority of EP neurons in the posterolateral motor territory are firing tonically at relatively fast discharge rates, ventrolateral (VL; rodent equivalent of Vop) connecting neurons are largely held in a burst ‘ready’ mode. However, in pathologically dystonic rats, under the same recording conditions, in which the firing activity of EP neurons is appreciably slower (and more irregular and bursty), VL neurons are largely in an erroneous tonic mode. From these experimental observations, in the resting state, GPi appears be chiefly controlling the burst versus tonic firing mode, rather than involved in direct linear inhibition of the VL neurons. Thalamocortical relay neurons are known to have the intrinsic ability to generate bursts in response to inhibitory synaptic inputs. This has been observed in GABAergic brainstem inputs to reticular thalamic neurons in rodents (Bokor et al. 2005). The normal shift from the tonic to burst mode of firing has been attributed to a threshold hyperpolarization setting of the basal resting membrane potential (Bokor et al. 2005; Mooney et al. 2004). This in turn, has been attributed to hyperpolarization-induced de-inactivation of voltage-gated T-type Ca^2+^ channels (Jahnsen and Llinás 1984; Jahnsen and Llinás 1984; Deschenes et al. 1984). Low threshold calcium current, upon depolarizing input, results in a long-lasting low threshold spike (LTS) crowned by a burst of Na^+^/K^+^ spikes. From our experimental observations in normal and dystonic rats, we postulated that the spatial and temporal synaptic summation of inhibitory inputs from EP neurons controls the strength and duration of net inhibitory postsynaptic potentials (IPSP) and, in turn, the hyperpolarization state of VL neurons.

This study simulates the role of different firing patterns observed in rat EP under normal and pathological conditions on a computational model of Vop neuron. The aim will be to determine the differential role of various input patterns in controlling the firing mode (tonic verses burst) and other firing properties (rate and irregularity) of the Vop model.

## 2. Material and Methods

Here we focus on two key factors in modeling GPi neuronal influence on the behavior of thalamic Vop neurons. These factors include firstly, modeling the burst and tonic behavior of Vop neurons and secondly, modeling the synaptic weights of multiple GPi inputs to Vop. Both of these factors are developed in depth in this section, along with the presentation of supportive experimental neuronal discharge behavior of EP and VL neurons in healthy and dystonic rats at rest.

### 2.1. Preliminary Experiments and Data Collection

To characterize thalamic motor-related neuronal firing properties and its primary pallidal influences, we collected and analyzed extracellular microelectrode recordings in VL and EP from normal and dystonic rats under awake, resting head restraint conditions. All experiments were approved and monitored by the Hunter Holmes McGuire Veterans Affairs Institutional Animal Care and Use Committee (IACUC) and performed in accordance with regulatory guidelines.

#### Animals

Normal, healthy Wistar Gunn rats were used for the normal studies. Dystonia is humans and experimentally in rats is characterized by prolonged abnormal co-contractions of opposing and unintended muscle groups. For the dystonia studies, kernicterus (Chaniary et al. 2009) and GP (rodent equivalent of GP externa (GPe)) lesion (Kumbhare et al. 2017) rat models were used. The GP lesion model is induced by administering the fiber sparing neurotoxin ibotenic acid into a posterior ventrolateral hotspot region in the motor territory of GP, which induces contralateral dystonia with co-contractions of antagonist muscle pairs within 4-5 hrs of the injection. The animals display contorted truncal posturing and unilateral dystonic limb extension indistinguishable from that induced more generally in kernicterus rats.

#### Surgery

The surgical and recording procedures were previously described in detail (Kumbhare et al. 2015). Briefly, surgeries were carried out under 1.5-4% general isoflurane anesthesia (with 1 L/min O2) in 40-47-day old rats. A custom stainless-steel head fixture (Chaniary et al. 2011) was firmly secured to the animal’s skull using miniature screws and epoxy. The local anesthetic bupivacaine (0.1-0.55 ml) was injected into the incision and the analgesic buprenorphine (0.25-1.6 mg/kg, i.p.) was administered prior to discontinuing the isoflurane. After the initial surgery, the animal was returned to their cage and allowed 24 hrs for recovery. No overt signs of stress, pain, or change in behavior were evident after the surgery. The following day, the rat’s head was immobilized by clamping a custom stainless-steel head fixture into a custom stereotaxic positioner (Chaniary et al. 2009). Inhalation anesthesia (isofluorane 2–2.5%) was briefly delivered and a 3.5 mm burr hole (2 mm caudal, 1.5 mm lateral to the bregma) was drilled into the skull exposing the underlying duramater. After 30 min, allowing for full recovery from effects of anesthesia, neuronal recording sessions were initiated.

#### Recording

An Eckhorn microelectrode manipulator (Thomas RECORDING GmbH, Giese, Germany) was mounted onto a KOPF stereotactic arm and the relevant motor territories of EP or VL were targeted based based on rat brain atlas coordinates (Paxinos and Watson 2013). Extracellular neuronal activity was recorded using high impedance (1–2 MΩ), 100 μm Thomas RECORDING quartz-platinum microelectrodes, for 60–120 s at a sampling rate of 40–44 kHz, and amplified and passed through a band-pass filter (gain=50, bandwidth 0.07–8 kHz) via an AlphaLab SnR data acquisition system (Alpha Omega Co. USA Inc., Alpharetta, GA, USA).

#### Data processing and analysis

The data were filtered, spike sorted, and the spike trains were discriminated as described previously (Kumbhare et al. 2015).

### 2.2. Computational Model for VL neuron

All computational modelling and simulations were performed in MATLAB Simulink 9.1.

#### 2.2.1. Modified Hodgkin and Huxley model

A modified (simplified) Hodgkin and Huxley model of a typical thalamic neuron was generated that includes fast sodium (Na^+^), potassium (K^+^), leakage, and T-type calcium (Ca^2+^) channels (Rubin and Terman 2004). The derivative of the membrane potential of the thalamic neuron (v_th_) can be described in terms of the various channel currents as given by

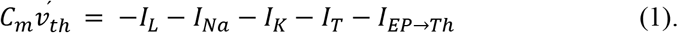

Here C_m_ is the membrane capacitance. I_EP→Th_ is the cumulative synaptic input currents from the various EP neurons. I_L_, I_Na_, I_K_, I_T_ are the leakage, sodium, potassium, and T-type calcium current respectively. These voltage-dependent currents are formulated by following equations:

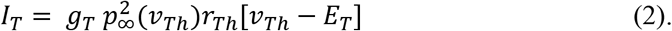

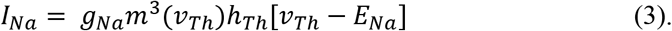

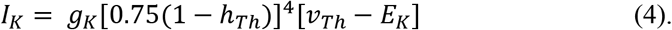

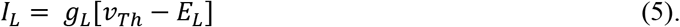

The membrane conductance (g_T_, g_Na_, g_K_ and g_L_) and equilibrium reverse potentials (E_T_, E_Na_, E_K_ and E_L_) are described in Table 1. The other terms are described by the voltage dependent dynamic equations 6 through 15 below as follows:

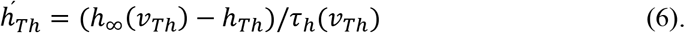

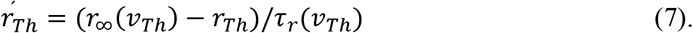

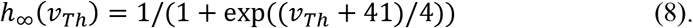

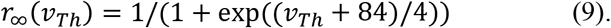

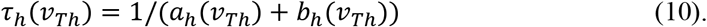

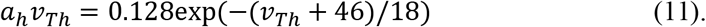

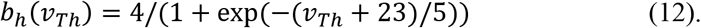

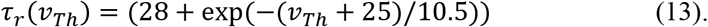

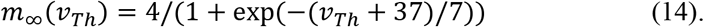

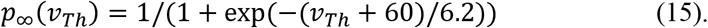

**Table 1.**
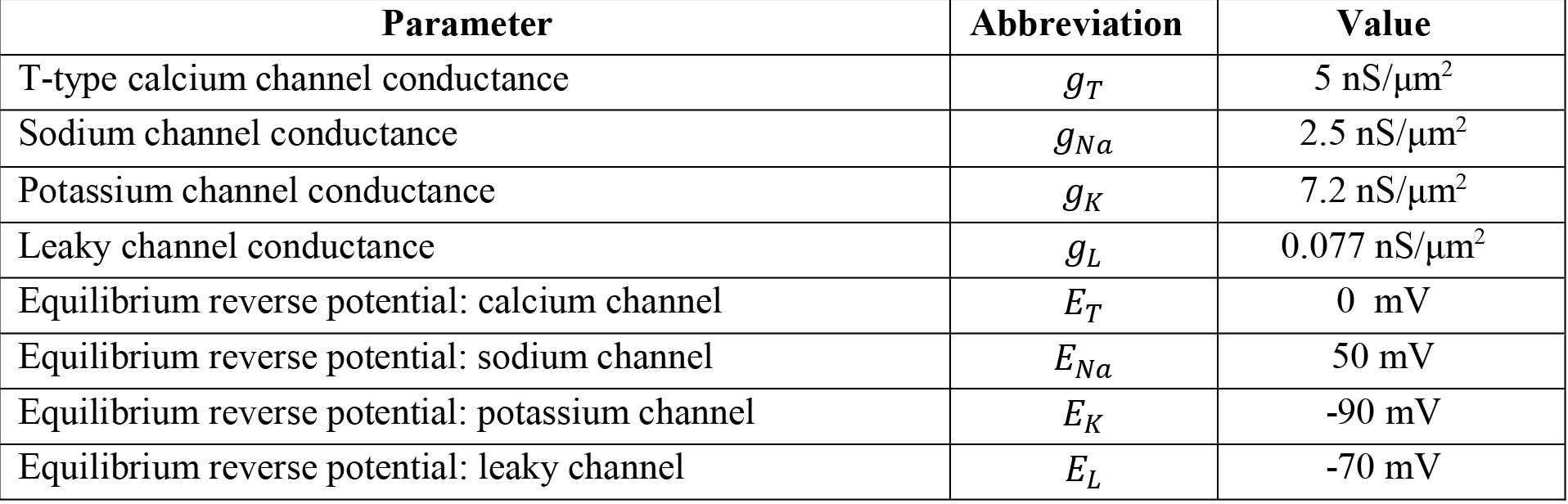
Parameter used to model the thalamic VL neuron.

| Parameter | Abbreviation | Value |
| --- | --- | --- |
| T-type calcium channel conductance | $g_T$ | 5 nS/$\mu\text{m}^2$ |
| Sodium channel conductance | $g_{Na}$ | 2.5 nS/$\mu\text{m}^2$ |
| Potassium channel conductance | $g_K$ | 7.2 nS/$\mu\text{m}^2$ |
| Leaky channel conductance | $g_L$ | 0.077 nS/$\mu\text{m}^2$ |
| Equilibrium reverse potential: calcium channel | $E_T$ | 0 mV |
| Equilibrium reverse potential: sodium channel | $E_{Na}$ | 50 mV |
| Equilibrium reverse potential: potassium channel | $E_K$ | -90 mV |
| Equilibrium reverse potential: leaky channel | $E_L$ | -70 mV |

The corresponding MATLAB Simulink block diagram is included in the Appendix (A1-A4).

#### 2.2.2. Depolarization/ pace-making drive

The depolarization of the membrane potential to initiate neuronal firing could be triggered either by self-pacing ability (Ramirez et al. 2004; Prinz et al. 2003) of the neuron itself or by external excitatory inputs (Frère and Lüthi 2004; Kiehn et al. 2000; Szucs et al. 2003). The intrinsic ability of a neuron to generate action potentials results from an interplay between the dynamics and densities of different ion channels (Ramirez et al. 2004; Blankenship and Feller 2010). Both ion channel density and activity dependent ion channel dynamics (i.e. membrane potential as a function of ionic concentration) modulates the ionic conductivities (Zhang and Linden 2003; MacLean et al. 2003). For a fixed combination of ionic channels, a wide range of discharge patterns could be obtained by varying the conductance of the ionic channels (Prinz et al. 2003). A spontaneous pace-making can be achieved by shifting the balance of ionic and leakage channels in such a way that it creates conditions of net inward current at resting potential. A higher conductance of sodium channel can facilitate spontaneous self-paced depolarization of the membrane potential, by initiating influx of current. The higher conductance of potassium, on the other hand, can control the spontaneous firing rate by controlling the rate of repolarization.

In the present model, to simulate intrinsic pace-making behavior in thalamic relay neurons (Deschenes et al. 1984), including pallidal receiving neurons (Destexhe and Sejnowski 2009; Yu et al. 2004), the conductance of Na^+^ and K^+^ was increased intermittently (within the acceptable range), while keeping the leakage conductance constant. Simulated injection of an external excitatory current increases the firing frequency of the neuron (by speeding up the depolarization) while an inhibitory current decreases the discharge frequency. However, the inhibitory current decreases the frequency only down to a threshold level where it then hinders the pace-making activity altogether (by compensating the inward current due to leakage and ionic channel thus disrupting any ability of depolarization).

#### 2.2.3. Integration of synaptic inputs

A separate Simulink block was implemented to simulate the synaptic inputs onto Vop (appendix Fig.A6.a). The intention here was to assign variable input weights at synapses and to simulate synaptic conditions induced by incoming pallidal GABAergic spike trains. All the spike train inputs from GPi were differentially weighted prior to being summed. Synaptic weights of different input spike trains were assigned empirically in order to generate the output patterns observed for certain input types. In some Dystonic cases we had only nine inputs while other Dystonic and healthy cases had 10 inputs. Each spike (action potential) pulse induces release of GABA vesicles from the presynaptic GPi terminal. The released GABA molecules attach to the GABA postsynaptic receptors on Vop neurons, opening chloride ions channels. The Cl^−^ influx, in turn, generates a negative inward post-synaptic membrane current. An integrator, a low pass filter, in the Simulink block (appendix Fig.A6a) simulates this by converting the incoming discrete voltage pulses into a continuous input current. Also, additional simulated factors, including the concentration of Cl^−^ and GABA in the extracellular and intracellular fluids (ECF and ICF), the number of GABA receptors, and the GABA receptacle pump present on the transmitting (EP) axons all cause non-linear effects on the post-synaptic potentials (PSP). This dynamic response is modelled by a feedback loop in the Simulink block (appendix Fig.A6a) that adaptively modifies the decay factor for the combined IPSP. The shape of the feedback function (appendix Fig.A6b) determines the rate of decay (appendix Fig.A6c). A single excitatory input can also be added to the synaptic block, to depolarizes the VL neuron in addition to the self-pacing drive.

*Design of the feedback function* (appendix Fig.A6b): The incoming GABA spike generates a transient hyperpolarization or IPSP at the synapse, which decays towards the resting potential over time. If the spike trains are at slower tonic rate, the input synaptic current to VL will be low and the decay function will be similar to a Resistor–Capacitor circuit (RC circuit) with a fixed decay time constant. Faster input spike trains retain the hyperpolarized state for longer duration. The combined buildup of the IPSPs will overcome the decay rate. The flat portion of the feedback loop represents this constant decay rate. If fast input spiking continues, more Cl^−^ channels are recruited before the previous Cl^−^ channels can shut off, this will cause stronger IPSP build-up with an even slower decay rate. However, if even faster spiking is maintained for longer, the GABA receptors and the Cl^−^ and GABA concentration will reach saturation. This will limit further build-up of IPSP causing a steep increase in the decay rate. Thus, any further increase in neuronal firing rates in EP will have no effect on the postsynaptic potentials of VL neurons.

Transfer characteristics as well as the Simulink© model for the synapse block are presented in Fig.5A of the appendix. The synaptic weights are shown in Table A1 of the appendix and filter block is shown in the Appendix in Fig.5A.

### 2.3 Generation of EP input trains

After the model was tested in an isolated environment with direct input currents, it was subjected to multiple patterned simulated spike train synaptic inputs. Each set of spike trains was created based on simplified and realistic experimental observations from neuronal recordings in the EP of healthy and dystonic animals.

#### 2.3.1. Preliminary training data

The preliminary training data used a baseline average firing rate and firing mode observed in EP in normal rats (Figs.4 and 5). The regular spike trains were generated using equally spaced time stamps with an inter-spike interval equal to the reported mean firing rate in EP in rats (though mildly slower than that observed for GPi in primates (Vitek et al. 1998)). Burst trains as observed in EP with movement were generated by alternating high frequency time stamp epoch and equally spaced pauses.

#### 2.3.2. Physiologically observed spike train

For testing the model’s efficacy on physiological data sets, we simulated multiple spike trains based on the properties observed in normal and dystonic rats (Fig.1). The simulation techniques were detailed previously (Kumbhare and Baron 2015). Briefly, a neuronal spike train data pool was generated using complex spike train modeling with control over various discharge features. These features include refractory limitations, level of irregularities, burstiness, and temporal non-homogeneity. The baseline inter-spike sequences were derived from gamma probability distribution based on two free parameters, regulating both regularity level and firing rate. Inter-spike intervals were randomly chosen from this exponential distribution to generate the time stamps with an additional limitation of refractory period. For burst trains, each spike in the above sequence was replaced by a burst or a non-burst event, depending on the desired properties. Different combinations of these EP (representative GPi) input patterns were then fed in the Vop neuron model and the corresponding weights were calculated to produce the desired firing mode in Vop.

**Fig. 1.**
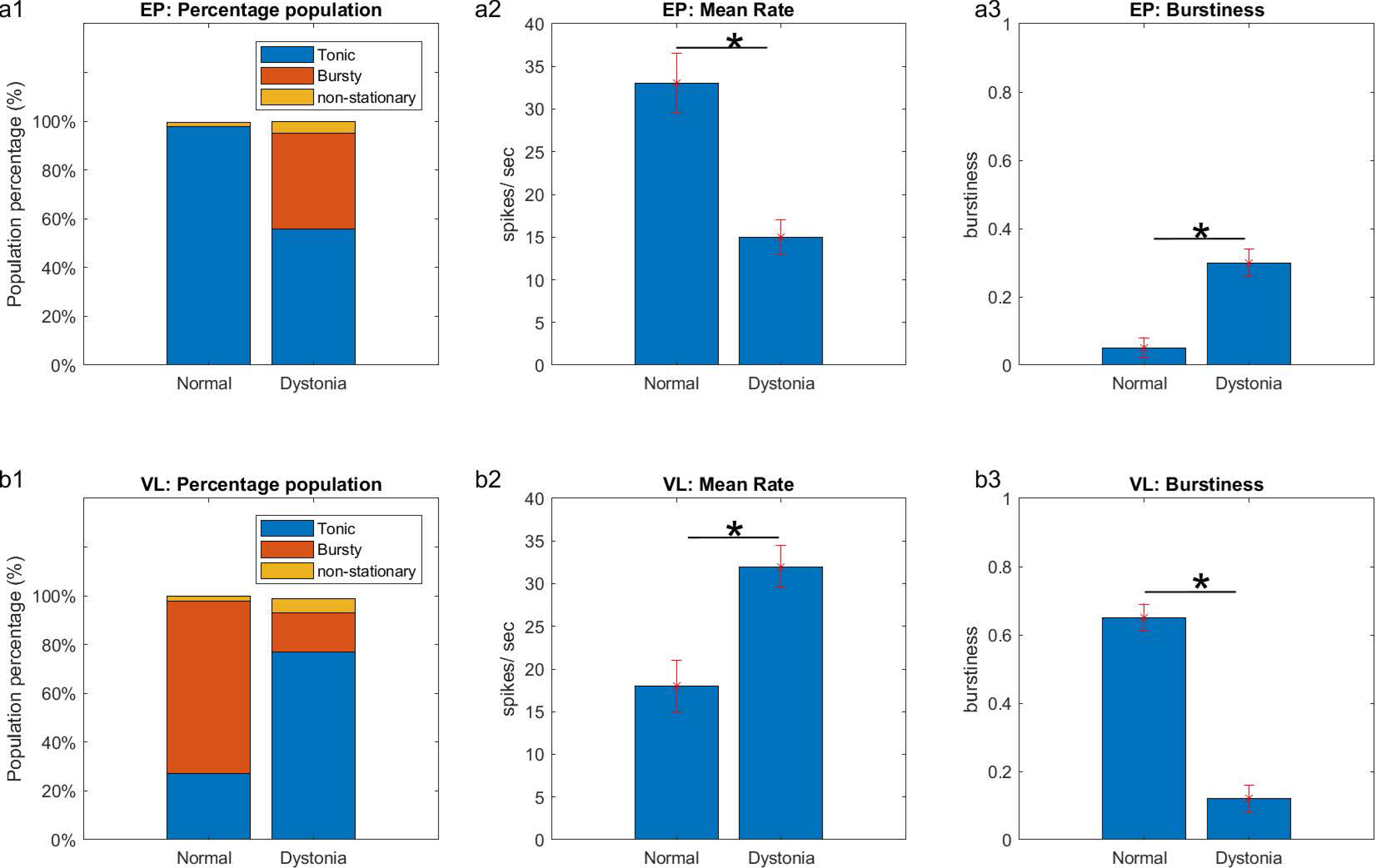
Characterization of neuronal patterns in different populations in entopeduncular nucleus (EP) and ventrolateral thalamus (VL) thalamus at rest in rats. Neuronal populations, mean firing rates, and mean burstiness distributions in EP (a1-a3) and VL (b1-b3) are shown for normal rats and for dystonic rats. Indicated are errors of the means. EP data was previously reported in different format (Kumbhare et al. 2015; Kumbhare et al. 2017). * indicates p < 0.05.

## 3. Results

### 3.1. Neuronal observations in EP and VL in normal and dystonic rats

Neuronal discharge activity was recorded from EP and VL in 27 normal rats and dystonic (both kernicterus and lesioned model) rats during awake rest (EMG silent) conditions. Our previous results (Kumbhare et al. 2015; Baron et al. 2011) revealed that during normal rest conditions, EP neurons (n=120) discharge for the most part (98%) in the tonic mode (Fig.1a), with tonic (either fast regular or slow irregular) discharge. Not previously reported, at rest, VL neurons (n=32) were observed to discharge at least 71% of the time in the burst (p≤0.05, one sample t-test; Fig.1b1). A percentage of recordings (2%) were unclassifiable and a percentage of single spikes represent LTS (and as such the burst mode) and would be incorrectly categorized as representing the tonic mode by our methodology. The overall mean firing rate of VL neurons was 18.2 ± spikes per sec and mean burstiness was 0.643 ± 0.12 (Figs.1b2, b3). Burstiness of the spike train is a burst metric based on burst parameters, including burst percentage, burst tendency, and burst entropy (Kumbhare and Baron 2015).

Previously (Kumbhare et al. 2015), we demonstrated that the resting discharge activity of EP neurons is slower, more highly irregular, and bursty (Fig.1a1 for population distribution) in dystonic rats compared to normal rats [Required permission will be obtained from the respective Journals]. In a subset of dystonic rats (n = 4), we also simultaneously recorded in VL. Not previously reported, in distinction from normal rats (p ≤ 0.05, one sample t-test), VL neurons (n=21) in dystonic rats predominately discharged in a tonic mode (77% classified as tonic vs 16% bursty). The total overall firing rate was increased by 77% to 32 ± 14.1 spikes/sec (p ≤ 0.05, one sample t-test) and burstiness was appreciably reduced to 0.12 ± 0.07 (p ≤ 0.05, one sample t-test) compared to that of VL neurons in normal rats (Figs.1b2 and b3).

### 3.2. Simulated contribution of different ionic channels on Vop mode

With a low level of external inhibitory current, the simulated thalamic neuron displays tonic output, as shown in the left part (see 0-400 ms) of Fig.2a. Pure excitatory inputs also keep the thalamic neuron in the tonic output mode, except when the neuron is driven at a threshold firing frequency higher than the refractory period (not shown). Lower levels of gradually increasing external inhibitory currents progressively decrease the tonic firing frequency. Larger values of external inhibitory currents completely silence the output of the neuron (see 400-650 ms, Fig.2a), consistent with in vitro observations (Jhansen and Llinas, 1983). The hyperpolarization activates Ca^2+^ channels permitting Ca^2+^ influx, which along with other factors, including interruption in external inhibitory input current, internal pace-making drive, and/ or external excitatory current input can lead to depolarization of the membrane potential. When the neuron transitions to the depolarized state from the maintained hyperpolarized state, it generates a burst pattern as shown in Fig.2a, between 650 −1000 ms; Fig.2 b,c). The simulated results illustrated in Fig.2 are consistent with those in literature (Suzuki and Rogawski 1989; Leresche and Lambert 2018; Rogawski 2002) and thus support the utility of our modeling of the LTS neuron to investigate the influence of different firing patterns of EP/GPi neurons on Vop neurons.

**Fig. 2.**
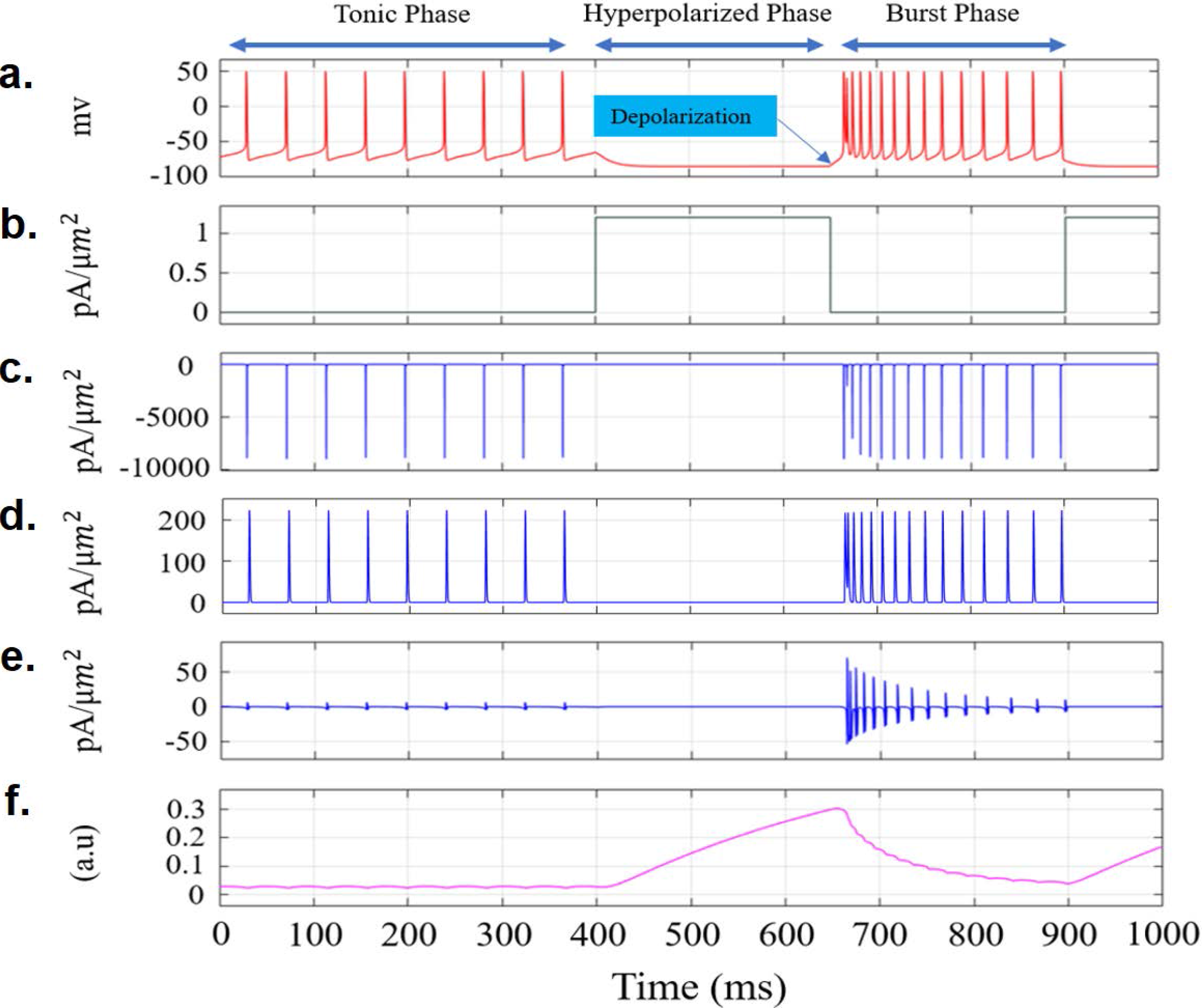
Contribution of different current channels during tonic, hyperpolarized, and burst phases of firing of the Vop neuron. **a.** Vop model output. External inhibitory current injection (b) causes the neuron to go to the hyperpolarized state. **b.** Magnitude of external inhibitory current profile. **c, d.** For sodium (c) and potassium (d) currents, upon inhibitory current injection, the magnitude and duration of their currents (i.e, spike potentials) do not change significantly. **e.** For calcium currents, a significant change is observed. **f.** The high magnitude of the current induced during the depolarization stage followed by the hyperpolarized state is explained by the rise in magnitude of inactivation variable *r*_*Th*_ of the calcium channel during prolonged hyperpolarization. Parameters used (Rubin and Terman 2004) are summarized in Table 1.

### 3.3. Behavior of Vop model with simplified simulated inputs

The Vop model was next subjected to simplified spike train inputs. These steps tested the influences of different EP/GPi firing patterns on the output of Vop model. Tonic trains were presented as spikes placed at regular intervals with slight randomness. Bursts were represented by transient high clustered activity followed by pauses at regular intervals. The primary goal of these inputs is to represent the input-output relationship in terms of baseline firing modes (tonic vs burst) observed in healthy and pathological (dystonic) conditions. Other properties of input discharge patterns (e.g., levels of irregularity and rate effects) were ignored in this simulation. The appropriate choice for synaptic weights for each GPi input is critical to the modeling. The weight distribution in simulated cases reflects quantitative involvement of specific input patterns (neuronal sub-populations) to produce the observed results.

#### 3.3.1. Healthy scenarios

As observed from in vivo recordings, there is an approximately equal (50/50) dichotomy of neuronal firing patterns in rat EP in the healthy state: 1) fast regular and 2) slow irregular patterns. Using the computational model of the Vop LTS neuron, we sought to understand the differential involvement of these two baseline input patterns in controlling the firing modes of Vop neurons. The fast discharging inhibitory input current (Fig.3c green box) places the membrane potentials of Vop neurons in a hyperpolarized state (Fig.3d, green box). If these fast inputs are continuously maintained, the hyperpolarized state will be sustained indefinitely, rendering the neuron incapable of discharging. For burst discharge, the neuron’s membrane potential which requires an overall reduction in the inhibitory inputs or an excitatory input (Fig.3c, red box). This will activate T-type Ca^2+^ channels, generating the Ca^2+^ crown for LTS burst (Fig.3d, red box). Thus, intermittent reductions in fast inhibitory inputs or excitatory cortical input, for example, is necessary for generation of LTS burst outputs. Fig.3 demonstrates this requirement in a healthy resting case. The regular tonic input patterns (Fig.3a, input 1 to 5) can maintain the hyperpolarized membrane state necessary for LTS but cannot produce the desired action potentials in the Vop model. The illustrated interrupted input (Fig.3a, input 6), on the other hand, allows the Vop model to reduce its hyperpolarized level and activate LTS burst generation. The high inter-pause rate in input 6 represents the collective effect of multiple inhibitory inputs to the neuron. *Thus, continuous fast inputs maintain the hyperpolarization and interrupted irregular inputs generate the depolarization*.

**Fig. 3.**
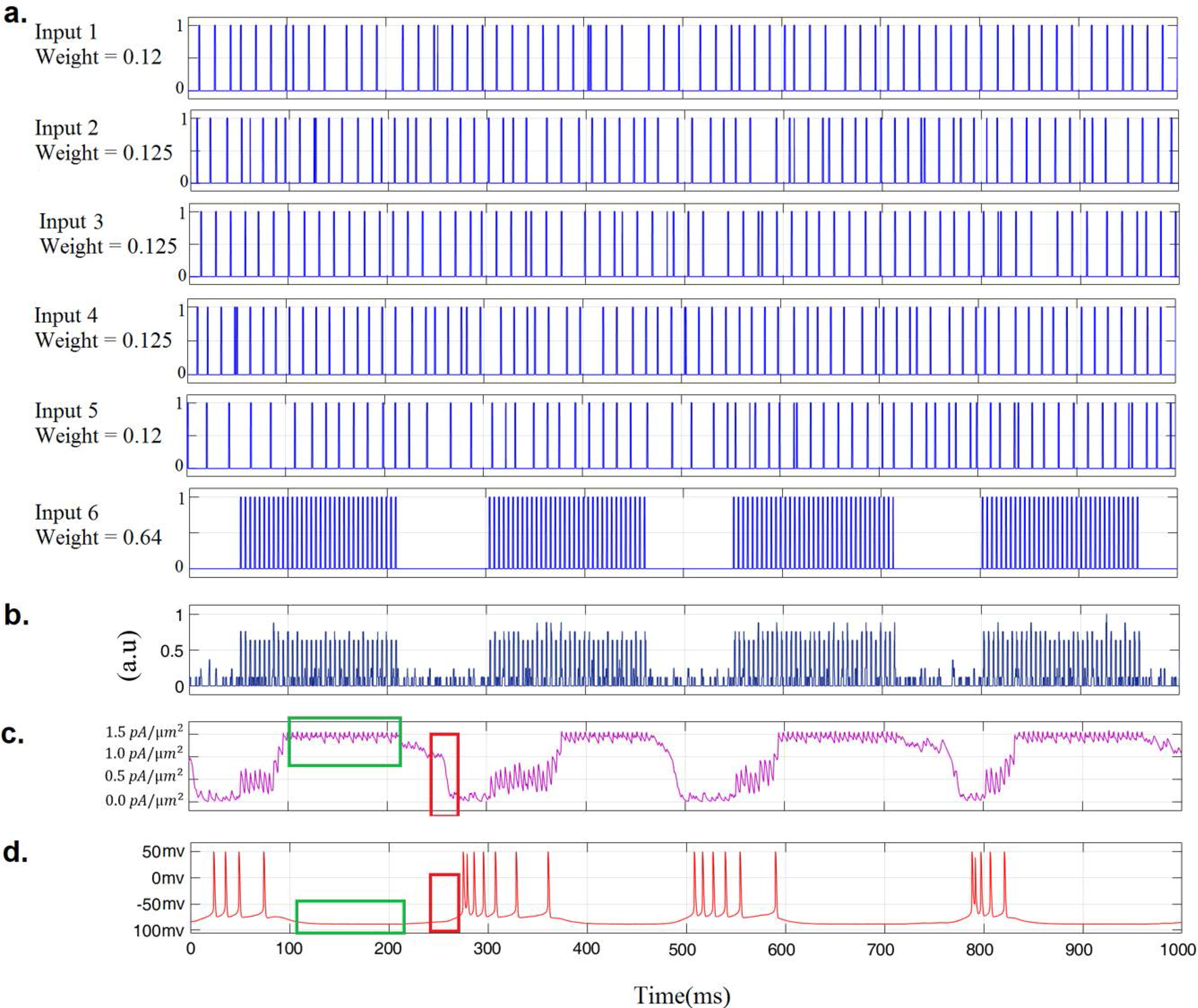
Vop model behavior under simplified simulated inputs, healthy scenario **a.** Simulated tonic inputs (Input 1-5) and an interrupted high frequency input (Input 6). **b.** Weighted sum of inputs 1-6. **c.** Synaptic block output after passing through the low pass filter. Green box shows saturated inhibitory current due to increased activity followed by a red box indicating decrease in inhibitory current due to lower activity. **d.** Resultant *bursty* output of the modeled Vop neuron. Green box shows a prolonged hyperpolarization followed by repolarization (red box) as activity level in the inputs decreases which ultimately leads to bursts.

#### 3.3.2. Dystonic scenario

A dichotomy in firing patterns in EP is preserved in dystonic rats, but is heralded by 1) highly slow and irregular and 2) slow bursty patterns. To mimic this behavior, three of the simulated simplified inputs were designated bursty (Inputs 1, 2 & 3) and the additional 3 (Inputs 4, 5 & 6) were set as tonic pattern, as shown in Fig.4. The overall reduction of firing rate in dystonia results in reduced inhibition, which had minimal effect on the baseline tonic firing mode of the VL neuron. Varying levels of simulated GPi tonic inhibitory output however alters the temporal spike discharges. Fig.4 demonstrates the behavior of the VL neuron model under dystonia-like EP inputs. The synaptic block output (Fig.4c) is reduced from that in than the healthy case (Fig.3c) indicating inadequate inhibition. This also results in inadequate hyperpolarization of the Vop neuron, reducing the possibility of bursts. With simulated burst inputs (Fig.4 a, inputs 1-3), the high frequency intra-burst spikes cause a transient increase in inhibition, but this is not sufficient to activate Ca^2+^ channel for burst generation. Since the burst inputs are more likely to have transient effects on the firing behavior, the weights of the burst inputs were increased. The stronger burst epochs (Fig.4a, red box) however are able to modulate the discharge properties of the output spike train (Fig.4d, red box). *In summary, while the pathological slow irregular and burst patterns modulate the baseline firing of Vop model neurons, they do not change the baseline tonic firing mode. In other words, in the case of reduced pallidal inhibition, GPi is unable to maintain sufficient hyperpolarization to place Vop neurons in the desired burst mode*.

**Fig. 4.**
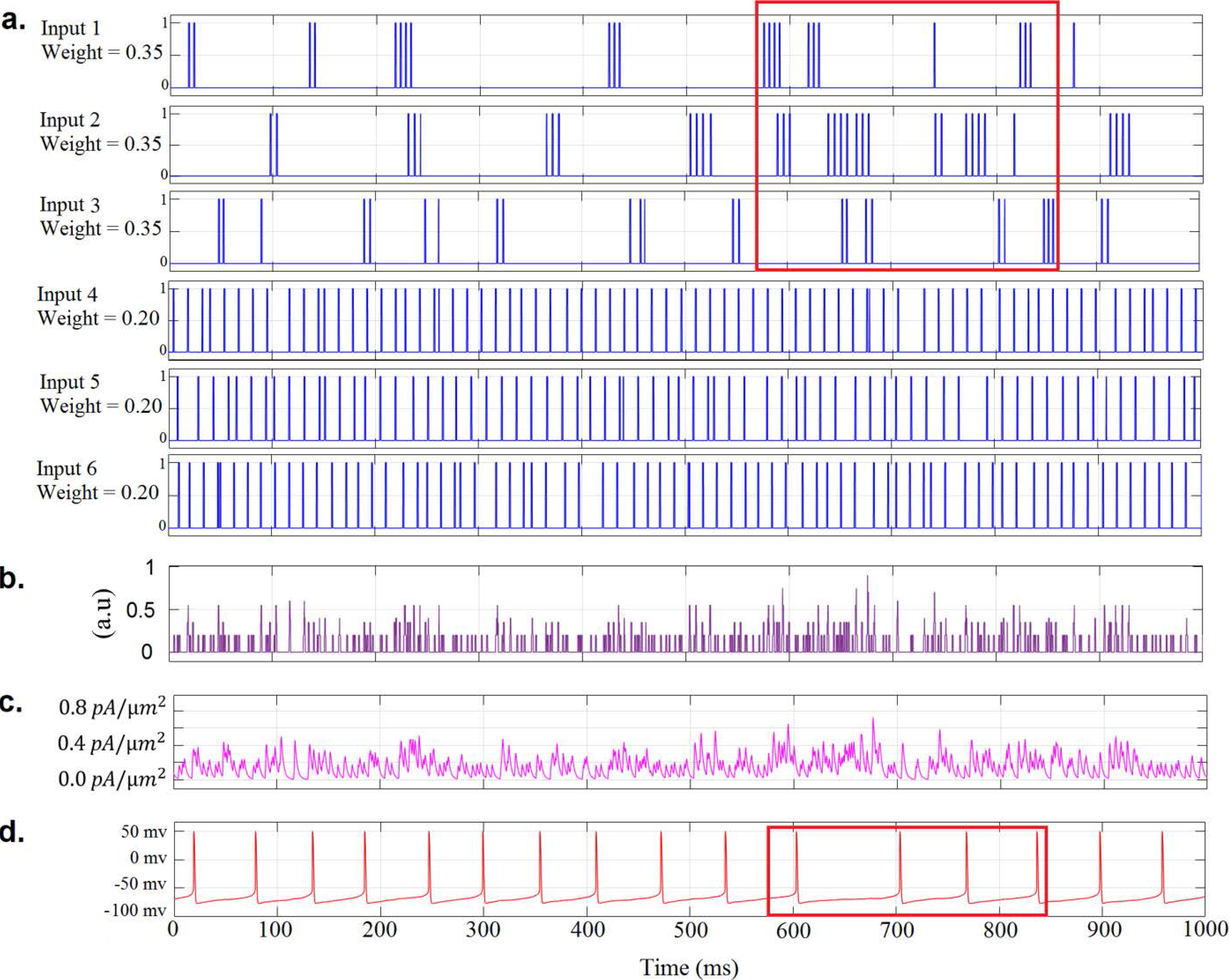
Vop model behavior under simplified simulated inputs, dystonic scenario **a.** Simulated bursty *(Input 1-3) and tonic inputs (Inputs 4-6)* Red box indicates more than usual overall activity due to closely packed bursts of different inputs. **b.** Weighted sum of inputs 1-6. **c.** Synaptic block output after passing through the low pass filter. **d.** Resultant *tonic* input of the modeled Vop neuron. A reduction in overall activity is indicated by the red box which corresponds to the increased activity indicated by red box in Fig.4.A.

If the burst events from different input trains fire in phase or in-sync with each other, the overall inhibitory activity can cause a hyperpolarization in Vop neurons and generate a burst. This occurs in advance of movement, when additional motor-related cortical and pallidal inputs drive multiple GPi neurons to fire in phase with the movement signals. The movement events however are outside the scope of the current manuscript.

### 3.4. Behavior of Vop model under physiological inputs

To further support the above arguments, and to test the efficacy of the model with real world scenarios, the model was subjected to representative EP neuronal inputs (Fig.5 and 6). The observed recordings in EP in healthy and dystonic rats were characterized and summarized previously (Kumbhare et al. 2017). Based on the summary, a pool of different input sets of representative 10 spike trains each were then simulated based on the representative firing patterns and properties of EP under healthy and dystonic conditions (Figs.5a, 6a and appendix Figs.A7a–10a).

**Fig. 5.**
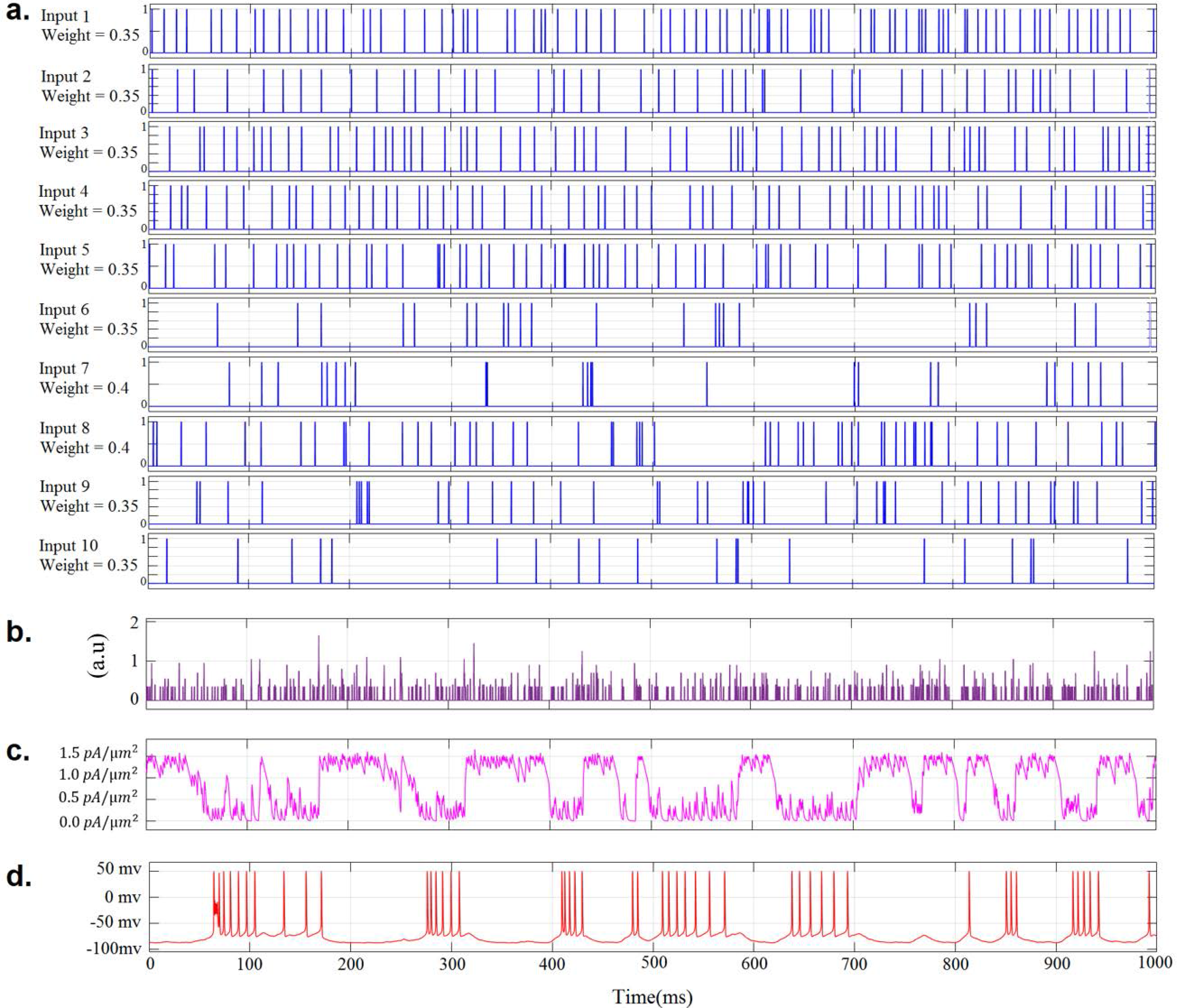
Vop model behavior under physiological inputs, healthy scenario *(details summarized under the case Normal 1 in Table II of supplementary document)*. **a.** Experimental pallidal inputs (*Inputs 1-10*) from healthy rats. **b.** Weighted sum of inputs 1-10. **c.** Synaptic block output after passing through the low pass filter. **d.** Resultant *bursty* output of the modeled Vop neuron.

**Fig. 6.**
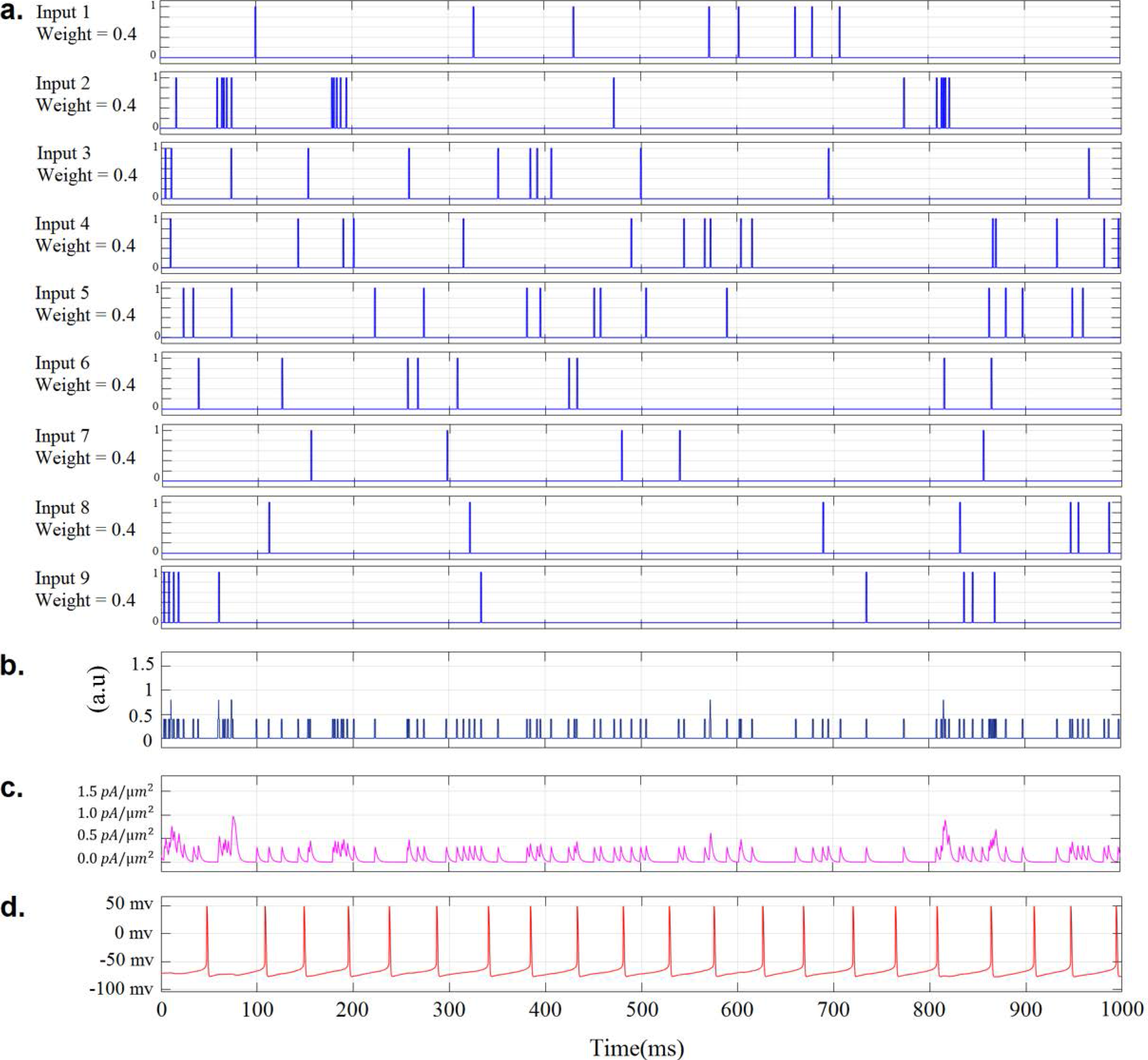
Vop model behavior under physiological inputs, dystonic scenario *(details summarized under the case Dystonic 2 in Table II of supplementary document)* **a.** Experimental pallidal inputs (labelled Inputs 1-9) from dystonic rats. **b.** Weighted sum of inputs 1-*9*. **c.** Synaptic block output after passing through the low pass filter. **d.** Resultant *tonic* output of the modeled Vop neuron.

#### 3.4.1. Healthy scenario

Fig.5 includes a simulated pool representing observed EP neuronal patterns of a healthy rat. Inputs 1-5 are fast regular tonic inputs (rate = 33.6 ± 6.2 spikes/sec; irregularity = 0.64 ± 0.04, and burstiness = 9.6 ± 6.6%) and inputs 6-10 are slow irregular inputs (rate = 17.0 ± 5.0 spikes/sec; irregularity = 1.27 ± 0.20, and burstiness = 16.0 ± 7.1%). The change in overall activity level of the EP neurons is indicated as the changes in the weighted summation of the EP inputs (Fig.5b). A total of four different sets of healthy rat senarios, with different combinations and properties of EP neuronal populations, were tested on the Vop model (Figs.5a, A7a, A8a). The firing rate, pattern, and weight distribution of the input train sets and the corresponding behavior of the output train from the Vop model are summarized in Table A1 of the appendix. It is apparent from the overall population’s coefficient of variation (CV) and burst percentage (BP) that the model works well for all of the natural inputs applied with the population ratio observed in EP of healthy rats.

To further understand the individual effects of each of the two types of healthy neuronal populations in EP, a set of only slow irregular spike trains (rate: 13.8 ± 5 spikes/sec) and only faster regular spike trains (41 ± 13) were used as inputs to the model Vop neuron (Table A1). Interestingly, with faster tonic inputs, the Vop neuron barely fires (mean firing rate = 4.1 spikes / sec). This suggests that the intermittent interruptions in the input inhibition to Vop are critical for maintaining the normal discharge rates of Vop neurons in the face of faster tonic input. In the case of purely slow irregular inputs to the modeled Vop neurons, the average firing rate increases, suggesting potentially higher discharge opportunities for the Vop neuron. On the other hand, with pure slow irregular input, the burst percentage is reduced by 65% compared to mixed population inputs. This indicates that the fast-regular inputs are essential for maintaining the hyperpolarized state necessary for burst outputs.

#### 3.4.2. Dystonia scenario

Fig.6 includes a similar simulated pool of 10 representative neurons in EP in dystonic rats. During an isolated burst, the activity is high for an individual EP neuron, but, due to the appreciably slow inter-burst discharge frequency and higher irregularity level of the spike events, the possibility of the combined inhibitory inputs to produce a sustained hyperpolarization is very low. As a result, there is suppressed de-inactivation of the T-type Ca2+ channels (which requires the hyperpolarized state to be maintained for a minimum period) and largely tonic firing is observed. This is confirmed from the summation of the inputs (Fig.6b), as well as from the current output of the synaptic block (Fig.6c). Hence, in the dystonic case, the thalamic Vop neurons are not held at rest in the preferred movement signaling burst state. All six dystonic input sets that have been tested are summarized in Table A1 of the appendix. See Fig.6 and Appendix Figs.A9 and A10 for representative inputs and outputs for 3 dystonic EP neurons. Similar to the healthy scenario, a set of individual neuronal dystonic units (irregular and bursty) were individually subjected to the Vop neuronal model. Both set of patterns generated largely (< 1.8 %) non-bursty tonic outputs, indicating that the baseline behavior of the model does not vary appreciably with either type of dystonic inputs. The bursty inputs however do generate higher non-stationarity, indicating a high local variance.

## 4. Discussion

In this study, computational modeling was used to investigate the influence of GPi neurons on the firing patterns of motor-related pallidal receiving Vop thalamic neurons was in the normal and in dystonia, as a representative pathological movement condition. The presented simulations and modeling were derived from our experimental studies in healthy and dystonic rats (Baron et al. 2011, Kumbhare et al. 2015, Kumbhare et al. 2017). The derived pallidal-thalamic model explains the differential roles of various neuronal sub-populations of GPi in modulating the discharge patterns in Vop. This modeling depicts a previously unrecognized role for GPi as a firing mode selector of Vop neurons and a precise temporal controller of thalamocortical signaling.

To understand the differential influences of wide ranges in firing rates and patterns on the discharge properties of Vop neurons, we generated a computational model of a typical Vop relay neuron using modified Hodgkin and Huxley equations (Hodgkin and Huxley 1952) with physiological GPi inputs. This model accounts for the properties of different ion channels, including the calcium channels (Perez-Reyes 2003; Suzuki and Rogawski 1989; Yu et al. 2004) involved in the LTS burst, as well as the integration different types of synaptic inputs. The role of the thalamic calcium channels are to induce LTS when depolarization is preceded by a sustained hyperpolarized state (Jahnsen and Llinás 1984; Steriad 1984). The amount of time the hyperpolarized state needs to be maintained to create an LTS burst is dependent on the activation and inactivation gate kinetics of the T-type calcium channel. This can be controlled by the interrupted inhibitory current (i.e., from GPi) to the Vop model neuron, which allows subsequent hyperpolarization and depolarization of the membrane potential, leading to burst Ca^2+^ crown. In absence of any inhibitory current, the Ca^2+^ channels are inactive and an excitatory (e.g., corticothalamic) pulse generates more tonic spikes in the model neuron. These findings are consistent with in vitro and in vivo reports and thus support that our Simulink Vop neuronal model effectively reproduces the natural condition.

The pallidal inputs to the neuron model were derived from our experimental observations in the posterior motor territory of EP (GPi motor-equivalent) in rats. The synaptic weights of the model were optimized to deliver the expected output observed in Vop in vivo under different healthy and pathological conditions. The choice of synaptic weights of the EP/GPi inputs are the most critical aspect of this modeling. The distinct weights assigned to different synaptic inputs indicates the involvement of certain neuronal sub-population in the transmitted signals. Per our modelling, the continuous fast regular inputs, as observed in healthy rats, maintain the hyperpolarized state of the thalamic LTS neurons, limiting the overall information relayed to the cortex, thereby commanding resting/ minimal muscle activities. In distinction, irregular slow discharging pallidal neurons interrupt the Vop hyperpolarization to generate the burst firing. Thus, the irregular neurons in our modeling were assigned relatively more synaptic weights. The burst mode in the thalamus is considered to be the more efficient mode for detection and transmission of alterations in input signaling (Crick 1984; Sharman 1996 and Sherman 2001). The greater weighting of the irregular inputs in our modeling also reflects its critical role in regulating the temporal firing of Vop neurons. In the dystonic condition, the pathological GPi neurons lose their regularity and fast discharging properties, leading to more tonic Vop outputs. Although the pathologically slow irregular and bursty patterns can in part modulate the local discharging of Vop neurons, they cannot maintain the desired burst firing mode.

Unlike the dichotomous fast regular and slower irregular tonic firing EP in normal rodents (Kumbhare et al. 2015), GPi neurons in primates are reported to exhibit largely fast irregular spike activity (Vitek et al. 1998). Irrespective, the combination of fast and irregular GPi discharge activity in primates would be expected to have similar combined effects on Vop neurons as observed for VL in rats. Further, under movement disorder conditions, both rodents (Baron 2011; Kumbhare and Chaniary 2015) and primates (Starr et al. 2005; McClelland et al. 2016; Xiao et al. 2010) similarly show prominent reductions in pallidal firing rates. Per our modeling, this pathological reduction in the pallidal firing rate is the principal cause for our experimentally observed alteration in VL from a predominant burst to the tonic firing mode.

In summary, our neuronal simulations suggest a role for GPi input to Vop as a firing mode selector. Our Simulink model effectively reproduced normal and pathological states under resting conditions. With movement, in addition to transient movement related epochs in incoming EP inputs, Vop neurons additionally receive depolarizing cortical inputs. While the study here focused on neuronal rest activity, modeling of pallidal-thalamic movement related signaling will require experimental characterization of neuronal signals during active movement.

## Acknowledgement

The authors thank George Weistroffer, M.S. for technical assistance with the animal studies.

## Appendix 1. Simulink Model of the VL neuron

**Fig. A1.**
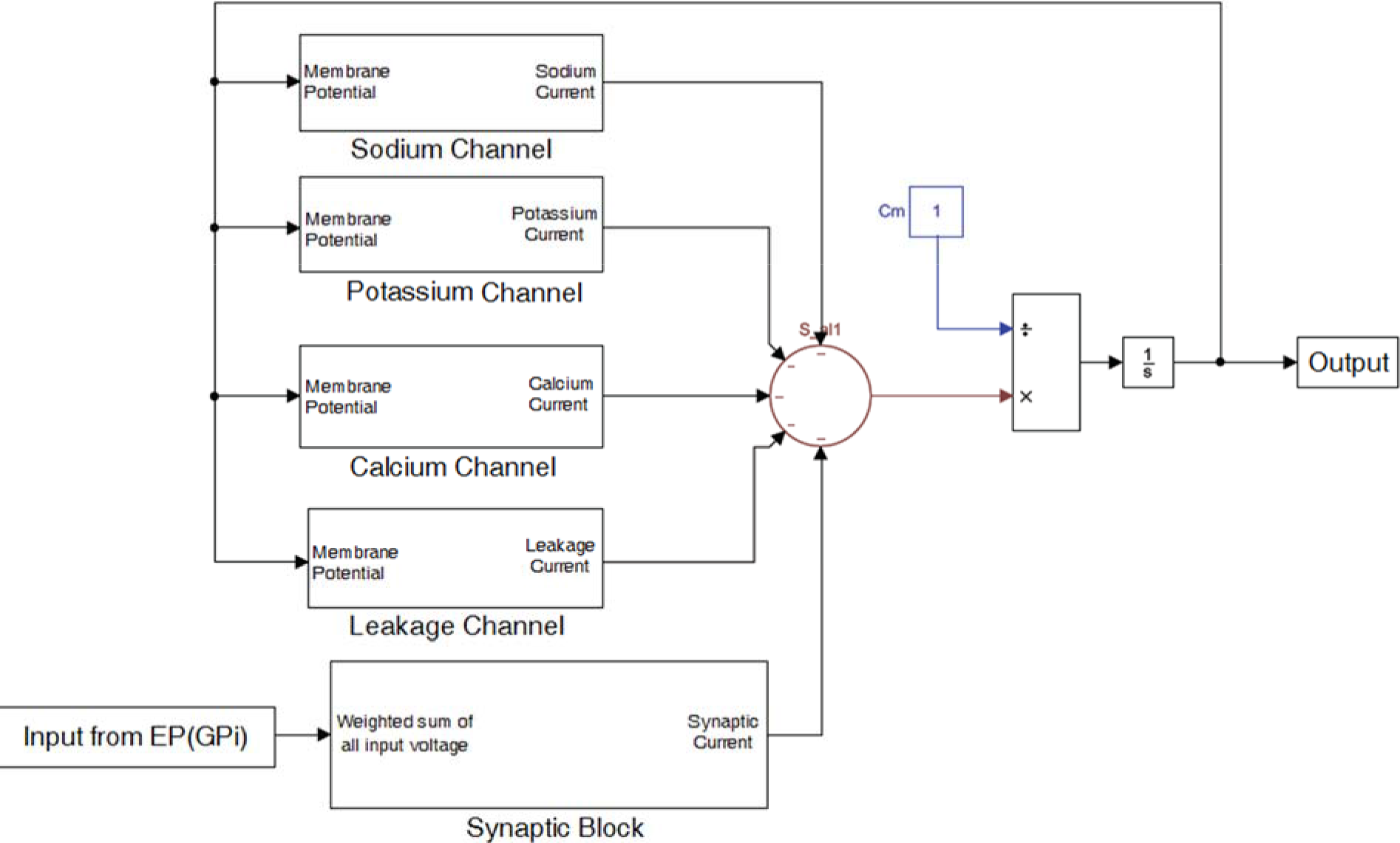
Simulink Model of the VL neuron with individual ion channels and synaptic block

**Fig. A2.**
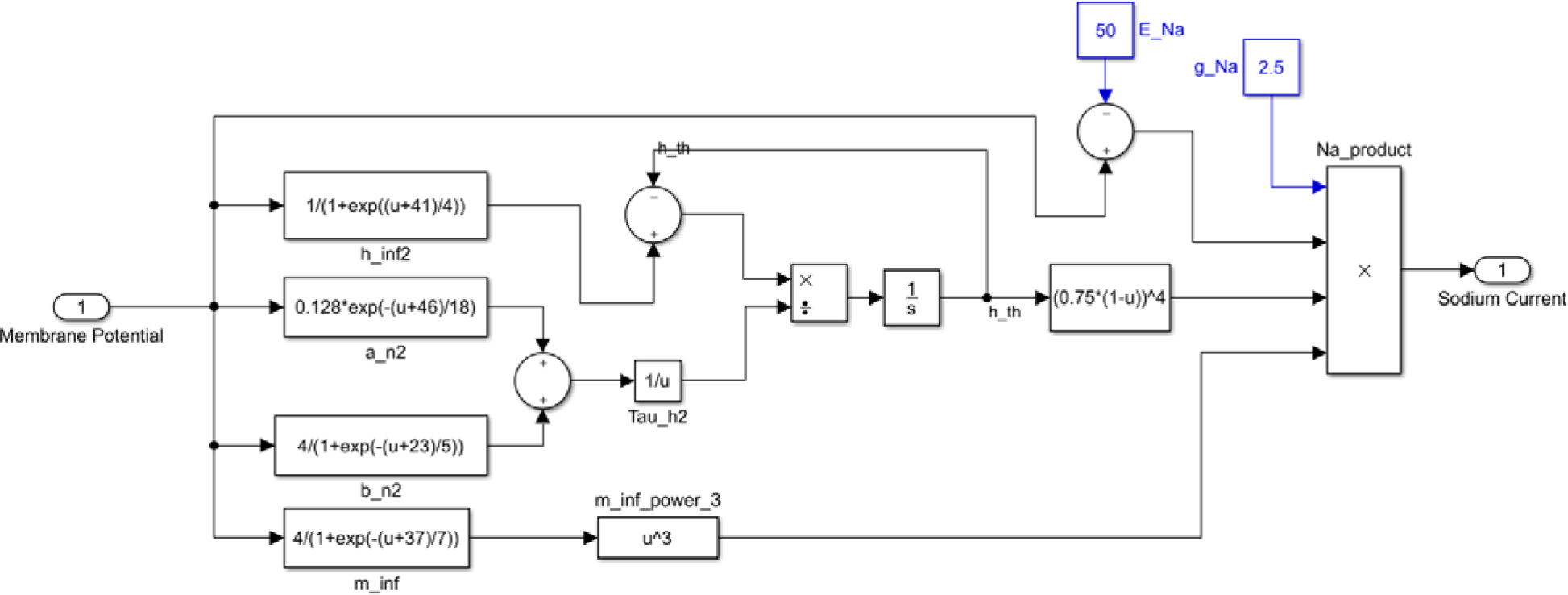
Detailed model of the Sodium Channel

**Fig. A3.**
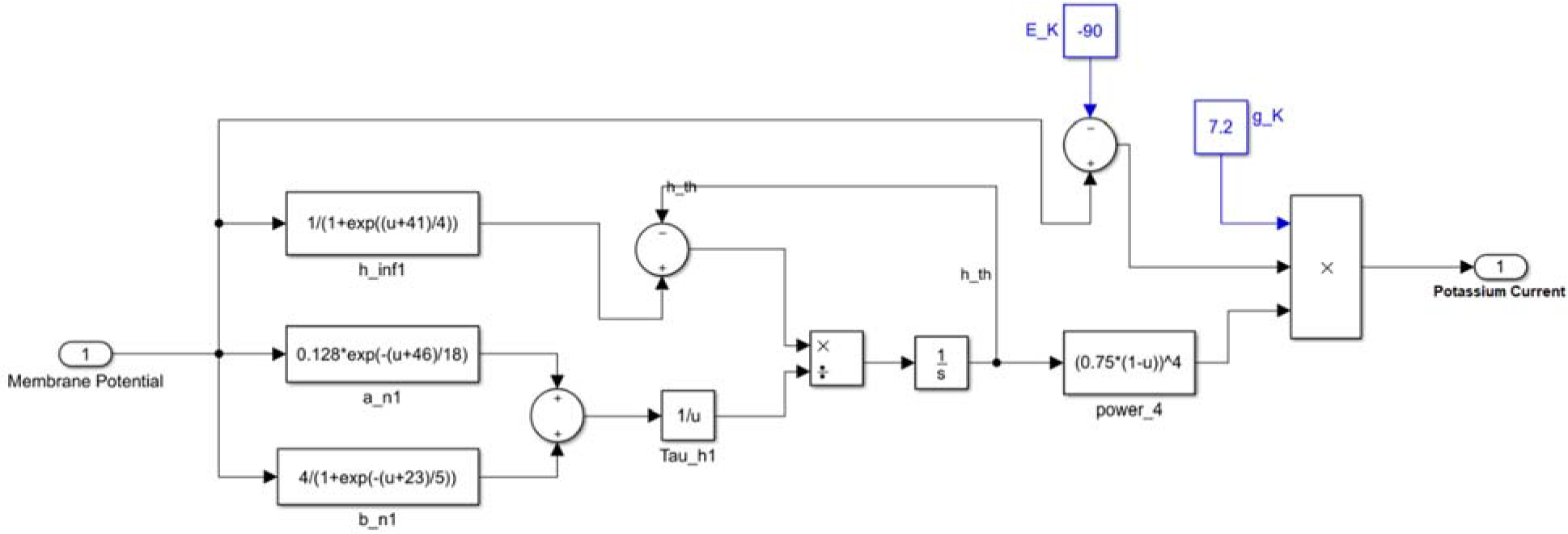
Detailed model of the Potassium Channel

**Fig. A4.**
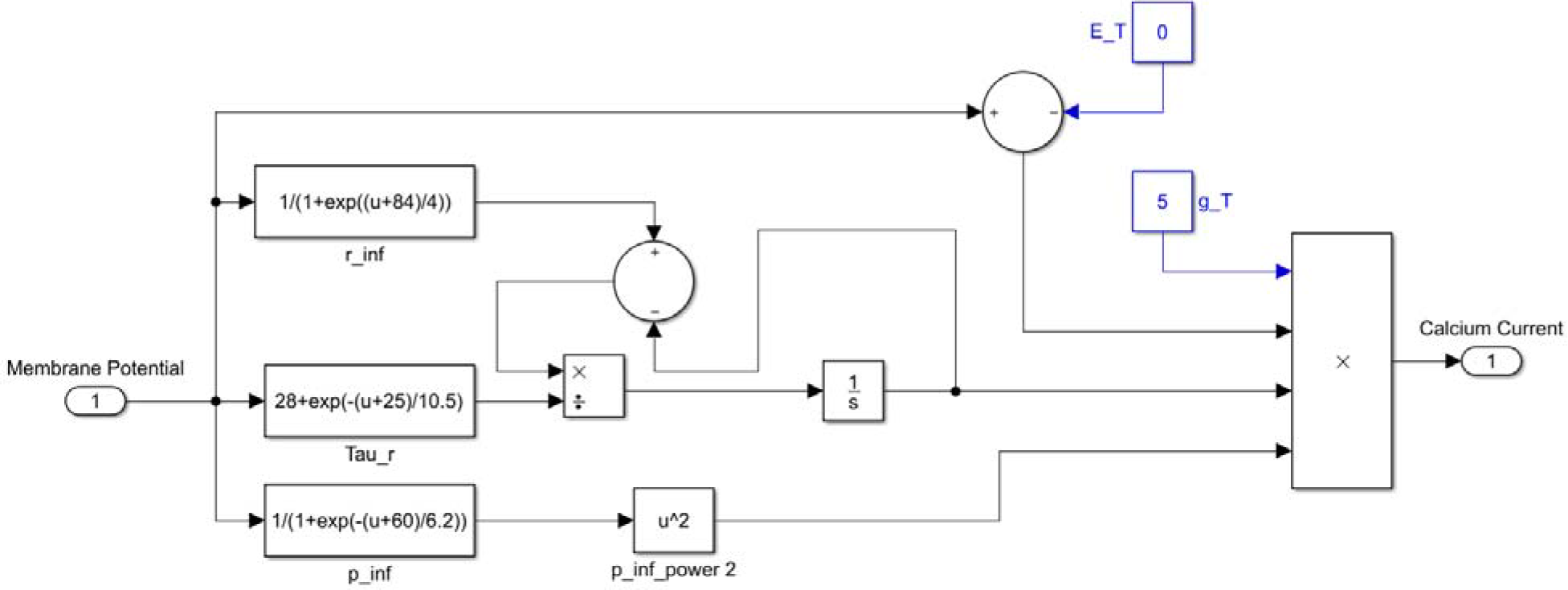
Detailed model of the Calcium Channel

**Fig. A5.**
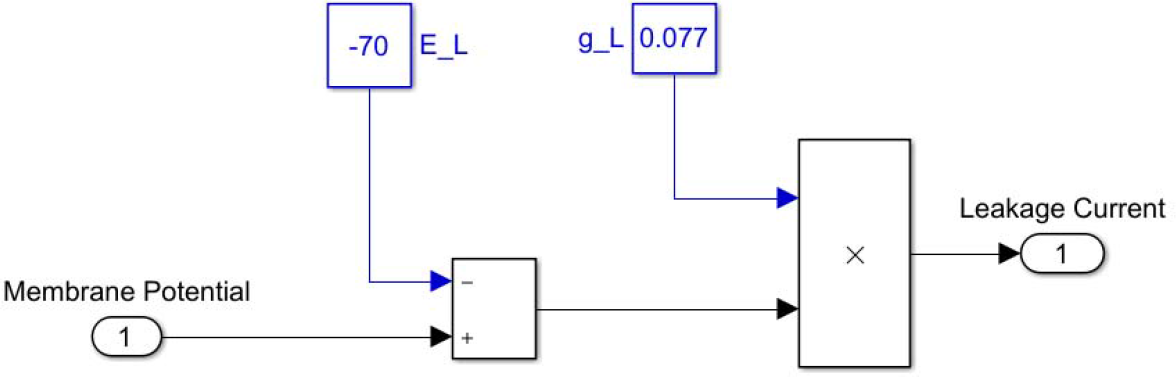
Detailed model of the Leakage Channel

## 2. The synaptic weights and synaptic block

**Fig. A6.**
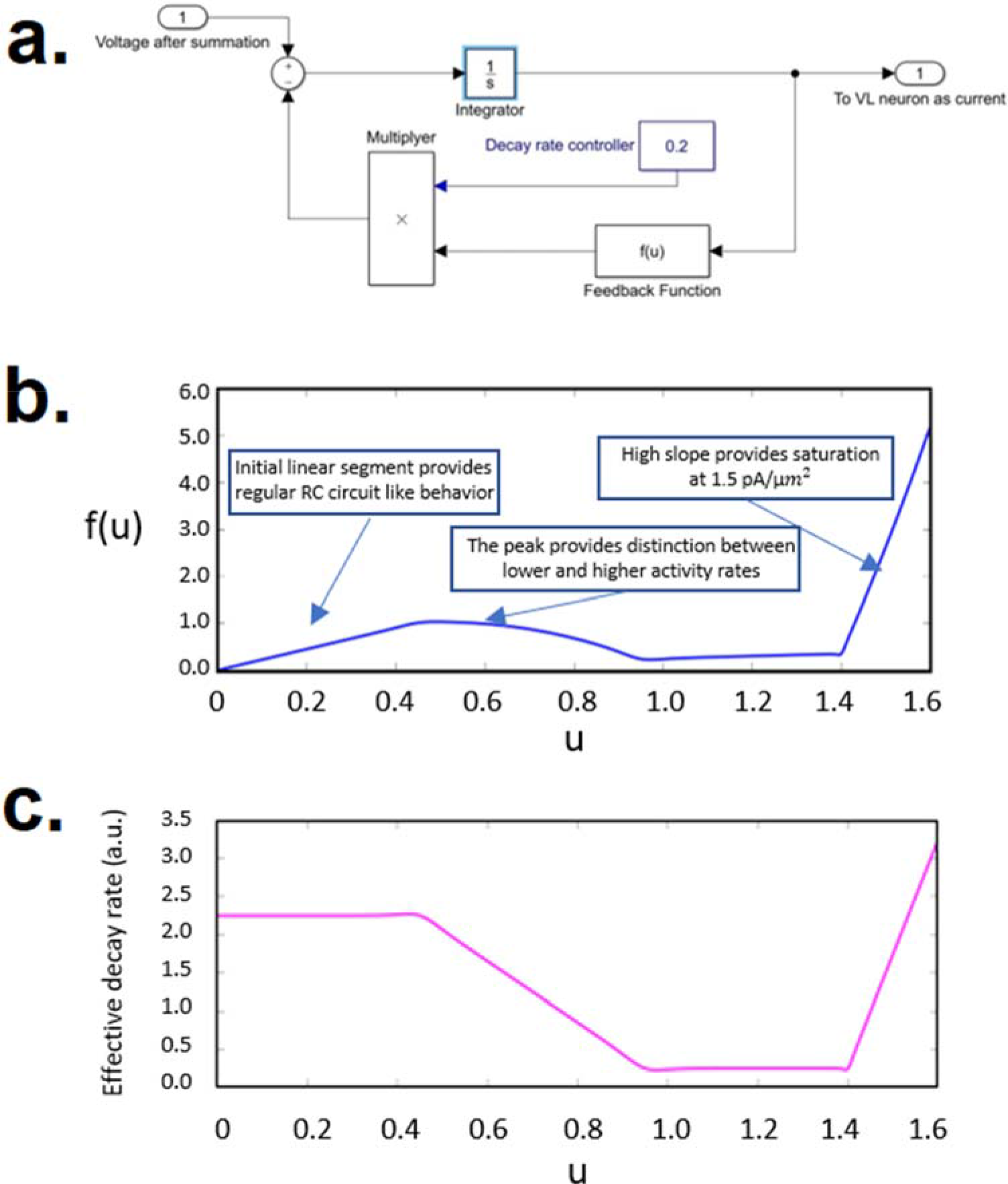
**a.** Simulink model of the synaptic block, **b.** “Feedback Function” output vs input providing different functionality as described in the plot, **c.** Thee effective decay rate of the synaptic block as described by the slope of the line connecting the origin and decay function for a given value of ‘u’

**Fig. A7.**
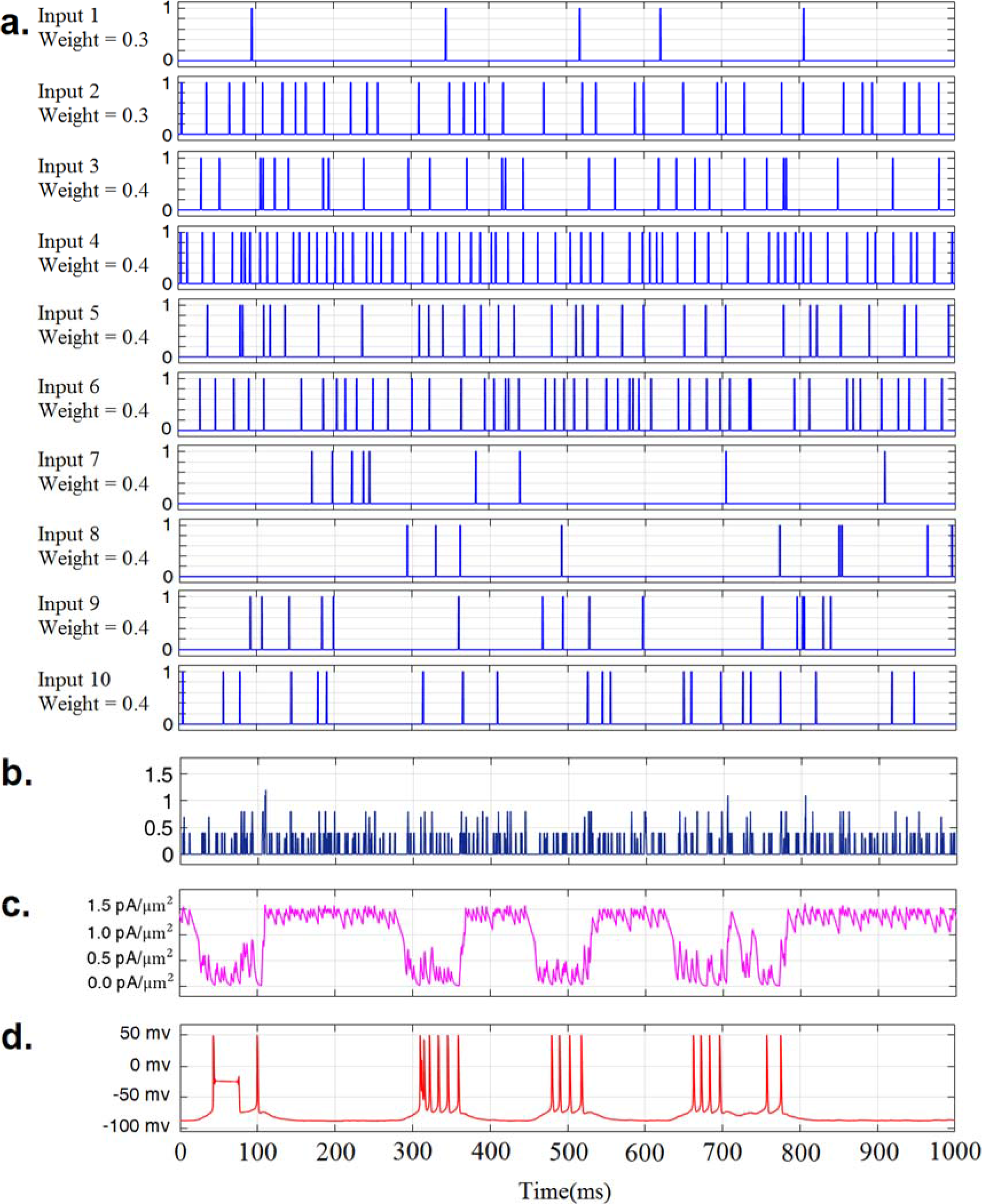
Vop model behavior under physiological inputs, healthy scenario *(details summarized under the case Normal 2 in Table II of supplementary document)*. **a.** Experimental pallidal inputs *(Inputs 1-10)* from healthy rats. **b.** Weighted sum of inputs 1-10. **c.** Synaptic block output after passing through the low pass filter. **d.** Resultant *bursty* output of the modeled Vop neuron.

**Fig. A8.**
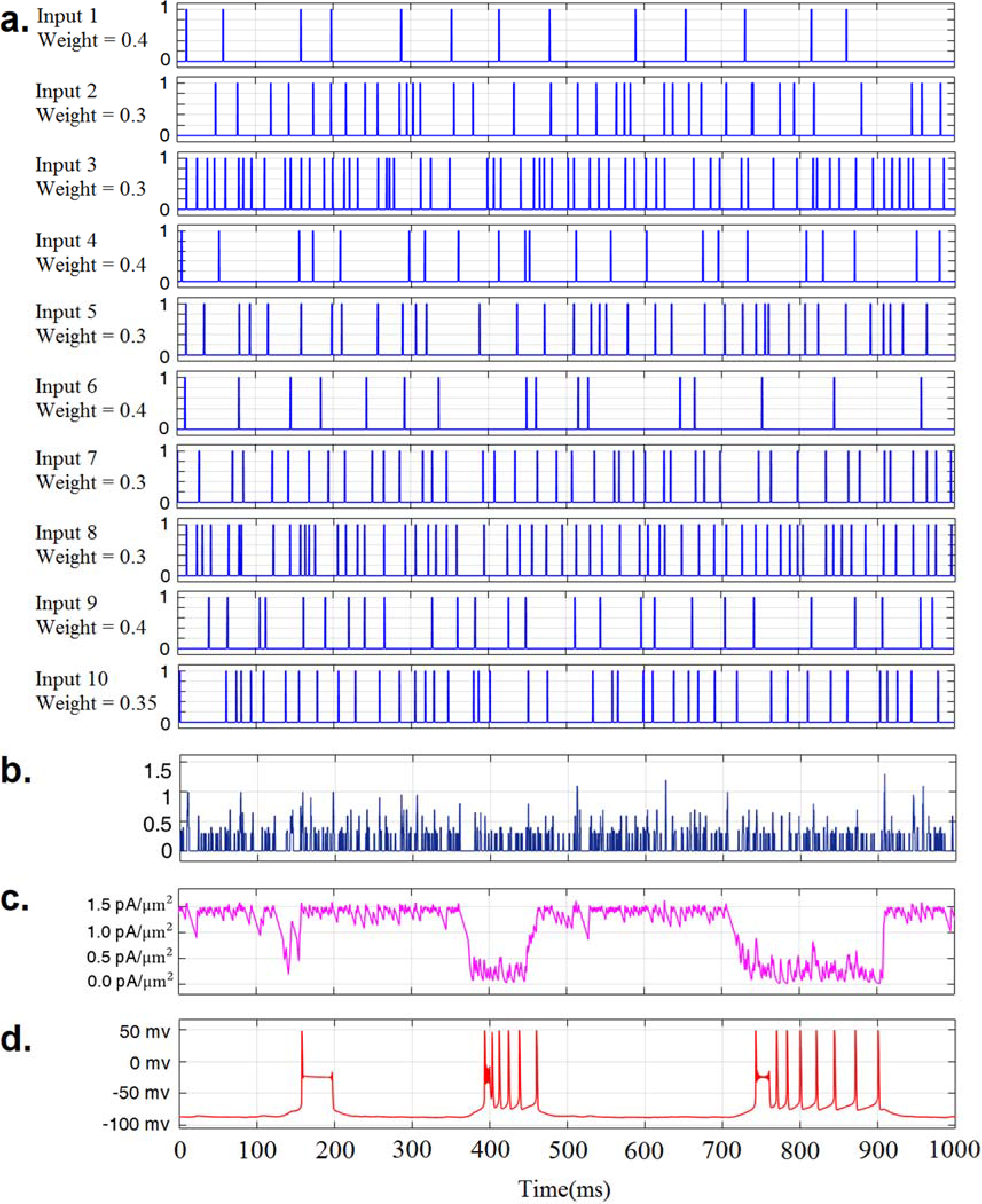
Vop model behavior under physiological inputs, healthy scenario *(details summarized under the case Normal 3 in Table II of supplementary document)*. **a.** Experimental pallidal inputs (*Inputs 1-10*) from healthy rats. **b.** Weighted sum of inputs 1-10. **c.** Synaptic block output after passing through the low pass filter. **d.** Resultant *bursty* output of the modeled Vop neuron.

**Fig. A9.**
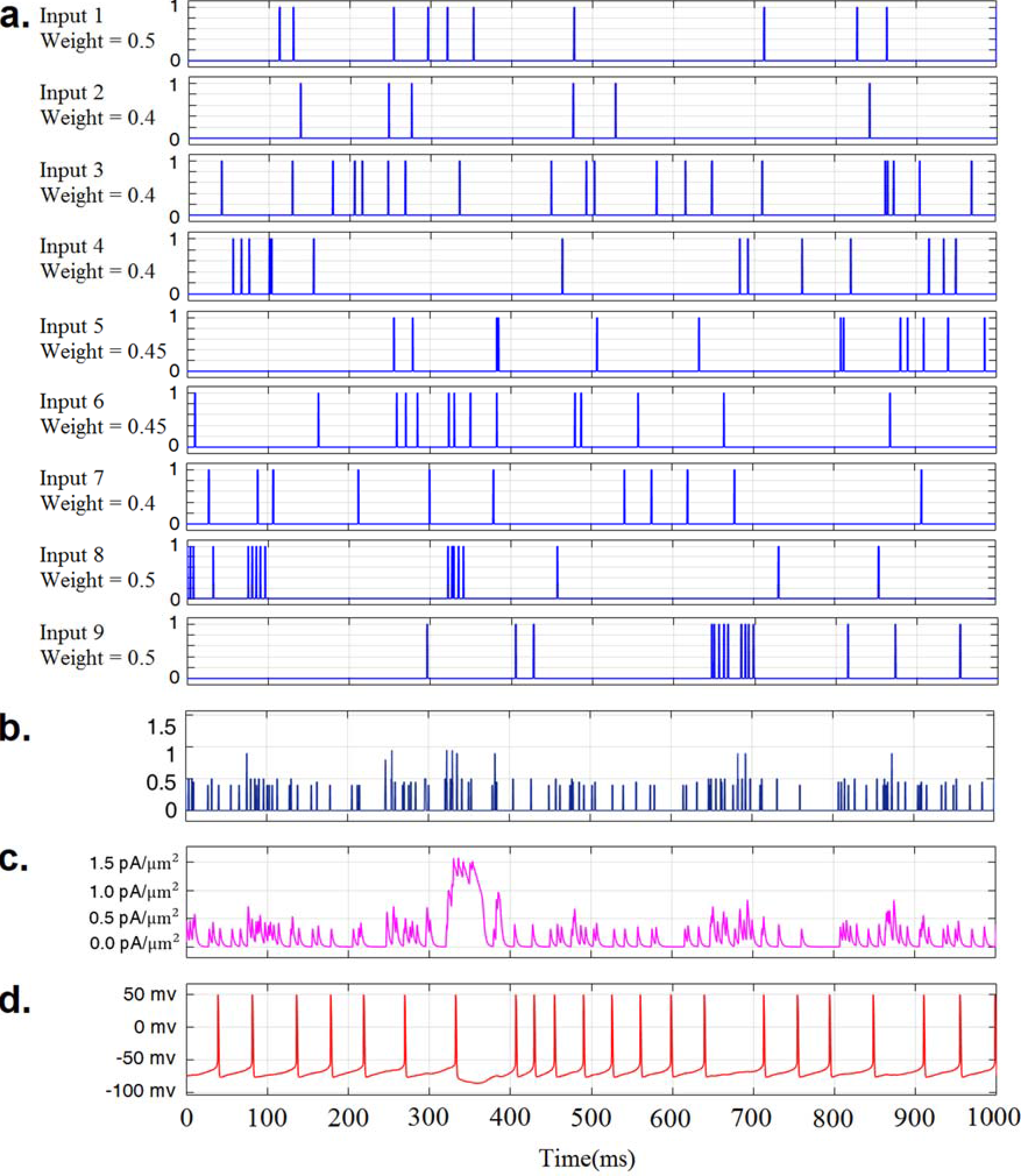
Vop model behavior under physiological inputs, dystonic scenario *(details summarized under the case Dystonic 1 in Table II of supplementary document)* **a.**Experimental pallidal inputs (labelled Inputs 1-9) from dystonic rats. **b.**Weighted sum of inputs 1-*9*. **c.**Synaptic block output after passing through the low pass filter. **d.** Resultant *tonic* output of the modeled Vop neuron.

**Fig. A10.**
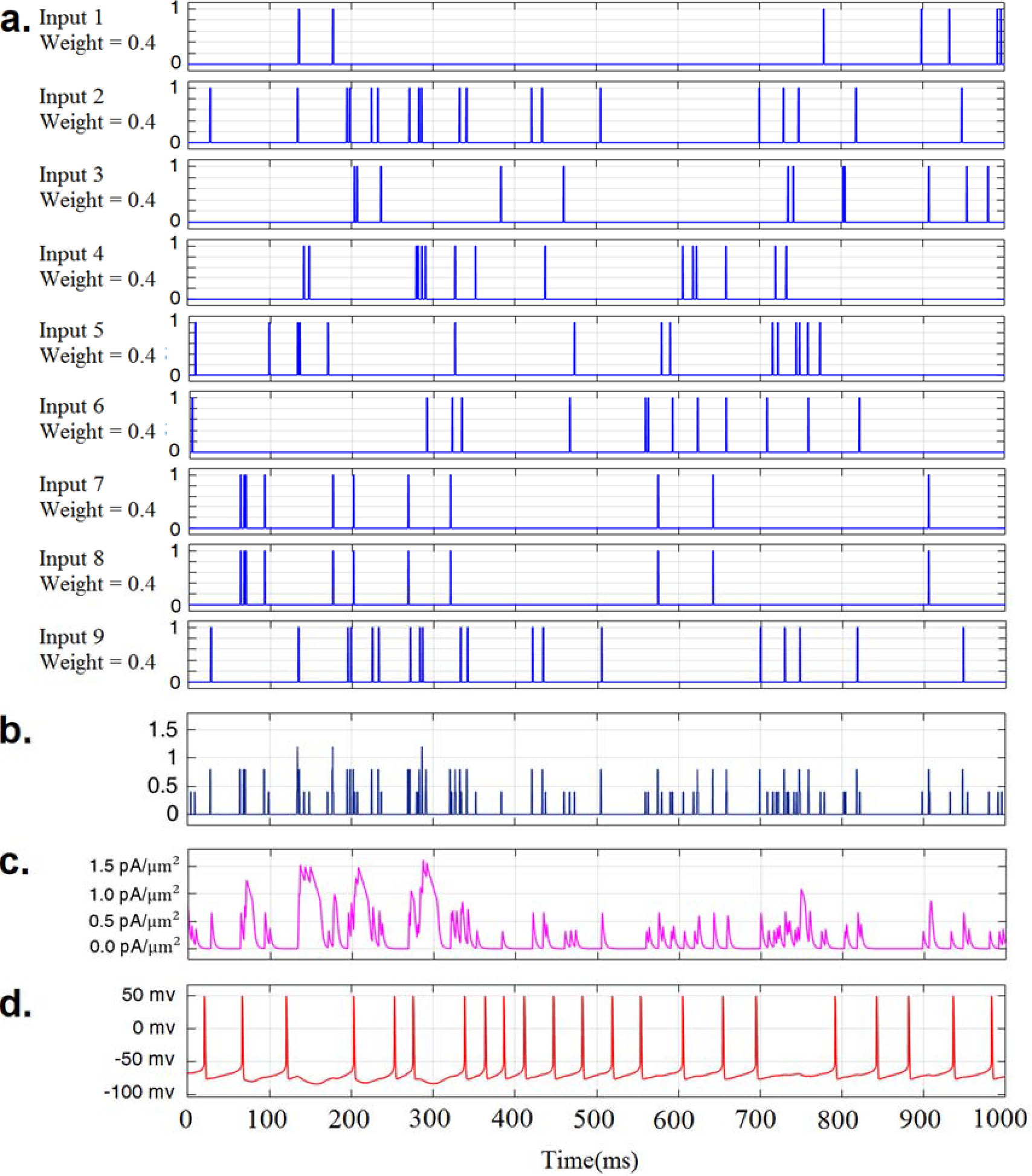
Vop model behavior under physiological inputs, dystonic scenario *(details summarized under the case Dystonic 3 in Table II of supplementary document)* **a.** Experimental pallidal inputs (labelled Inputs 1-9) from dystonic rats. **b.** Weighted sum of inputs 1-*9*. **c.** Synaptic block output after passing through the low pass filter. **d.** Resultant *tonic* output of the modeled Vop neuron.

**Table A1.** The firing rate, pattern, and weight distribution of the input spike trains and the corresponding pattern of the output train from the Vop model

| *Input set* | **Input/**<br>**output**<br>**number** | **Firing**<br>**Pattern** | **Firing Rate**<br>**(Spikes/**<br>**sec)** | **Coefficient**<br>**of Variation**<br>**(CV)** | **Burst**<br>**percentage**<br>**(BP) %** | **Assigned**<br>**synaptic**<br>**weights** |
| --- | --- | --- | --- | --- | --- | --- |
| *Normal*<br>*Set 1* | Input 1 | Irregular | 12.00 | 1.30 | 17.03 | 0.35 |
|  | Input 7 | Irregular | 14.72 | 1.26 | 19.01 | 0.4 |
|  | Input 8 | Irregular | 24.37 | 0.97 | 17.80 | 0.4 |
|  | Input 9 | Irregular | 19.67 | 1.07 | 22.41 | 0.35 |
|  | Input 10 | Irregular | 14.33 | 0.91 | 3.92 | 0.55 |
|  | Input 2 | Regular | 23.33 | 0.51 | 1.30 | 0.35 |
|  | Input 3 | Regular | 36.99 | 0.56 | 10.96 | 0.35 |
|  | Input 4 | Regular | 32.59 | 0.49 | 5.46 | 0.55 |
|  | Input 5 | Regular | 36.25 | 0.58 | 11.28 | 0.35 |
|  | Input 6 | Regular | 38.83 | 0.57 | 18.76 | 0.35 |
| | *Mean of*<br>*Input* | *Mixed* | $25.31 \pm 9.69$ | $0.82 \pm 0.30$ | $12.79 \pm 6.93$ | $0.40 \pm 0.08$ |
|  | **VL output** | **Bursty** | **14.00** | **1.73** | **25.21** | **NA** |
| *Normal*<br>*Set 2* | Input 1 | Irregular | 2.16 | 1.42 | 7.90 | 0.3 |
|  | Input 7 | Irregular | 8.57 | 1.14 | 7.45 | 0.4 |
|  | Input 8 | Irregular | 14.45 | 0.99 | 5.59 | 0.4 |
|  | Input 9 | Irregular | 11.22 | 1.44 | 16.93 | 0.4 |
|  | Input 10 | Irregular | 14.18 | 0.97 | 4.11 | 0.4 |
|  | Input 2 | Regular | 37.17 | 0.51 | 10.83 | 0.3 |
|  | Input 3 | Regular | 24.12 | 0.57 | 2.09 | 0.4 |
|  | Input 4 | Regular | 60.46 | 0.49 | 47.40 | 0.4 |
|  | Input 5 | Regular | 36.03 | 0.57 | 10.03 | 0.4 |
|  | Input 6 | Regular | 54.42 | 0.58 | 41.05 | 0.4 |
| | *Mean of*<br>*Input* | *Mixed* | $26.28 \pm 18.94$ | $0.87 \pm 0.36$ | $15.34 \pm 15.01$ | $0.38 \pm 0.04$ |
|  | **VL output** | **Bursty** | **10.60** | **2.10** | **21.14** | **NA** |
| *Normal*<br>*Set 3* | Input 1 | Regular | 14.06 | 0.52 | 0.00 | 0.4 |
|  | Input 2 | Regular | 35.58 | 0.57 | 12.09 | 0.3 |
|  | Input 3 | Regular | 41.09 | 0.57 | 21.71 | 0.3 |
|  | Input 4 | Regular | 57.75 | 0.51 | 47.19 | 0.4 |
|  | Input 5 | Regular | 22.76 | 0.57 | 2.61 | 0.3 |
|  | Input 6 | Regular | 36.17 | 0.54 | 12.23 | 0.4 |
|  | Input 7 | Regular | 14.77 | 0.50 | 0.00 | 0.3 |
|  | Input 8 | Regular | 40.79 | 0.56 | 18.34 | 0.3 |
|  | Input 9 | Regular | 56.01 | 0.48 | 39.96 | 0.4 |
|  | Input 10 | Regular | 23.04 | 0.57 | 1.96 | 0.35 |
| | *Mean of*<br>*Input* | *Regular* | $34.20 \pm 14.72$ | $0.54 \pm 0.03$ | $15.61 \pm 15.82$ | $0.35 \pm 0.05$ |
|  | **VL output** | **Slow and**<br>**less**<br>**bursty** | **4.13** | **1.55** | **10.87** | **NA** |
| *Normal*<br>*Set 4* | Input 1 | Irregular | 15.15 | 1.18 | 13.49 | 0.5 |
|  | Input 2 | Irregular | 4.10 | 0.92 | 0.00 | 0.4 |
|  | Input 3 | Irregular | 19.73 | 1.18 | 19.66 | 0.55 |
|  | Input 4 | Irregular | 10.18 | 0.94 | 4.25 | 0.4 |
|  | Input 5 | Irregular | 18.29 | 1.21 | 25.29 | 0.55 |
|  | Input 6 | Irregular | 15.43 | 1.25 | 22.01 | 0.5 |
|  | Input 7 | Irregular | 5.11 | 0.93 | 0.00 | 0.4 |
|  | Input 8 | Irregular | 20.93 | 1.12 | 19.62 | 0.6 |
|  | Input 9 | Irregular | 11.60 | 0.94 | 4.21 | 0.4 |
|  | Input 10 | Irregular | 17.58 | 1.05 | 12.31 | 0.55 |
|  | *Mean of Input* | *Irregular* | *13.81±5.59* | *1.07±0.12* | *12.08±8.97* | *0.49±0.07* |
|  | **VL output** | **Less bursty** | **23.598** | **0.81** | **8.23** | **NA** |
| *Dystonia Set 1* | Input 1 | Irregular | 11.008 | 1.106 | 7.212 | 0.5 |
|  | Input 3 | Irregular | 18.247 | 0.989 | 9.593 | 0.4 |
|  | Input 4 | Irregular | 15.562 | 1.072 | 15.809 | 0.4 |
|  | Input 5 | Irregular | 12.491 | 1.170 | 8.300 | 0.45 |
|  | Input 6 | Irregular | 12.168 | 1.114 | 9.362 | 0.45 |
|  | Input 2 | Bursty | 12.031 | 1.312 | 43.830 | 0.4 |
|  | Input7 | Bursty | 12.291 | 1.261 | 44.382 | 0.4 |
|  | Input 8 | Bursty | 10.632 | 1.306 | 46.502 | 0.5 |
|  | Input 9 | Bursty | 10.925 | 1.361 | 48.066 | 0.5 |
|  | *Mean of input* | *Mixed* | *12.82±2.36* | *1.19±0.12* | *25.90±17.89* | *0.44±0.04* |
|  | **VL Output** | **Tonic** | **21.74** | **0.37** | **0.81** | **NA** |
| *Dystonia Set 2* | Input 1 | Irregular | 12.25 | 1.13 | 12.500 | 0.4 |
|  | Input 3 | Irregular | 13.21 | 0.94 | 6.667 | 0.4 |
|  | Input 4 | Irregular | 15.05 | 1.087 | 8.014 | 0.4 |
|  | Input 5 | Irregular | 17.40 | 1.09 | 20.249 | 0.4 |
|  | Input 6 | Irregular | 6.33 | 1.09 | 4.032 | 0.4 |
|  | Input2 | Bursty | 14.06 | 1.40 | 51.210 | 0.4 |
|  | Input 7 | Bursty | 9.94 | 1.40 | 46.725 | 0.4 |
|  | Input 8 | Bursty | 12.85 | 1.45 | 48.276 | 0.4 |
|  | Input 9 | Bursty | 11.55 | 1.36 | 50.781 | 0.4 |
|  | *Mean of input* | *Mixed* | *12.52±2.97* | *1.22±0.17* | *27.61±19.85* | *0.4±0.00* |
|  | **VL Output** | **Tonic** | **21.53** | **0.35** | **0.00** | **NA** |
| *Dystonia Set 3* | Input 1 | Irregular | 7.59 | 1.21 | 9.93 | 0.4 |
|  | Input2 | Irregular | 14.94 | 1.10 | 12.85 | 0.4 |
|  | Input 3 | Irregular | 12.37 | 0.98 | 7.12 | 0.4 |
|  | Input 4 | Irregular | 15.58 | 1.11 | 15.41 | 0.4 |
|  | Input 5 | Irregular | 13.33 | 1.19 | 9.96 | 0.4 |
|  | Input 6 | Irregular | 9.52 | 1.17 | 7.14 | 0.4 |
|  | Input 7 | Irregular | 7.91 | 1.43 | 11.77 | 0.4 |
|  | Input 8 | Irregular | 12.35 | 0.92 | 7.12 | 0.4 |
|  | Input 9 | Irregular | 14.94 | 1.10 | 12.85 | 0.4 |
|  | *Mean of input* | *Irregular* | *12.06±2.88* | *1.13±0.14* | *10.46±2.82* | *0.4±0* |
|  | **VL Output** | **Tonic** | **21.48** | **0.36** | **0.00** | **NA** |
| *Dystonia Set 4* | Input 1 | Bursty | 14.09 | 1.23 | 48.28 | 0.4 |
|  | Input 2 | Bursty | 11.86 | 1.46 | 48.25 | 0.4 |
|  | Input 3 | Bursty | 13.80 | 1.18 | 39.63 | 0.4 |
|  | Input 4 | Bursty | 13.35 | 1.25 | 45.46 | 0.4 |
|  | Input 5 | Bursty | 12.26 | 1.25 | 46.53 | 0.3 |
|  | Input 6 | Bursty | 14.22 | 1.31 | 48.32 | 0.3 |
|  | Input 7 | Bursty | 11.37 | 1.24 | 45.65 | 0.4 |
|  | Input 8 | Bursty | 12.26 | 1.58 | 46.41 | 0.4 |
|  | Input 9 | Bursty | 13.92 | 1.46 | 47.16 | 0.4 |
|  | Input 10 | Bursty | 12.96 | 1.33 | 48.12 | 0.4 |
|  | *Mean of input* | *Bursty* | *13.01±0.97* | *1.33±0.12* | *46.38±2.48* | *0.38±0.04* |
|  | **VL Output** | **Tonic** | **22.55** | **0.51** | **1.87** | **NA** |
| *Dystonia Set 5* | Input 1 | Irregular | 9.66 | 1.25 | 13.95 | 0.5 |
|  | Input 3 | Irregular | 15.11 | 1.01 | 10.96 | 0.4 |
|  | Input 4 | Irregular | 11.26 | 1.29 | 19.28 | 0.4 |
|  | Input 5 | Irregular | 9.16 | 0.94 | 3.06 | 0.4 |
|  | Input 6 | Irregular | 12.17 | 1.25 | 17.99 | 0.5 |
|  | Input 7 | Irregular | 9.76 | 1.36 | 11.77 | 0.4 |
|  | Input 2 | Bursty | 12.99 | 1.40 | 50.20 | 0.3 |
|  | Input 8 | Bursty | 13.14 | 1.24 | 46.93 | 0.5 |
|  | Input 9 | Bursty | 15.11 | 1.47 | 50.41 | 0.3 |
|  | Input 10 | Bursty | 12.23 | 1.27 | 43.86 | 0.4 |
|  | *Mean of input* | *Mixed* | *12.06±2.02* | *1.25±0.15* | *26.84±17.72* | *0.41±0.07* |
|  | **VL Output** | **Tonic** | **21.24** | **0.33** | **0.00** | **NA** |
| *Dystonia Set 6* | Input 1 | Irregular | 2.37 | 1.24 | 0.00 | 0.5 |
|  | Input 3 | Irregular | 10.32 | 0.89 | 5.58 | 0.3 |
|  | Input 4 | Irregular | 7.51 | 0.99 | 7.84 | 0.4 |
|  | Input 5 | Irregular | 7.26 | 0.89 | 0.00 | 0.4 |
|  | Input 6 | Irregular | 21.80 | 1.18 | 26.16 | 0.3 |
|  | Input 7 | Irregular | 11.49 | 1.15 | 11.74 | 0.4 |
|  | Input 2 | Bursty | 12.57 | 1.36 | 45.26 | 0.4 |
|  | Input 8 | Bursty | 10.87 | 1.31 | 44.49 | 0.4 |
|  | Input 9 | Bursty | 13.17 | 1.18 | 47.44 | 0.5 |
|  | Input 10 | Bursty | 14.23 | 1.21 | 45.90 | 0.4 |
|  | *Mean of input* | *Mixed* | *11.16±4.85* | *1.14±0.16* | *23.44±19.51* | *0.40±0.06* |
|  | **VL Output** | **Tonic** | **21.74** | **0.26** | **0.00** | **NA** |

